# The Nucleosome Remodeling and Deacetylase complex has an asymmetric, dynamic, and modular architecture

**DOI:** 10.1101/2020.02.17.951822

**Authors:** Jason KK Low, Ana PG Silva, Mehdi Sharifi Tabar, Mario Torrado, Sarah R Webb, Benjamin L Parker, Maryam Sana, Callum Smits, Jason W Schmidberger, Lou Brillault, Matthew J Jackman, David C Williams, Gerd A. Blobel, Sandra B Hake, Nicholas E Shepherd, Michael J Landsberg, Joel P Mackay

## Abstract

The Nucleosome Remodeling and Deacetylase (NuRD) complex is essential for development in complex animals but has been refractory to biochemical analysis. We present the first integrated analysis of the architecture of the native mammalian NuRD complex, combining quantitative mass spectrometry, covalent cross-linking, protein biochemistry and electron microscopy. NuRD is built around a 2:2:4 pseudo-symmetric deacetylase module comprising MTA, HDAC and RBBP subunits. This module interacts asymmetrically with a remodeling module comprising one copy each of MBD, GATAD2 and CHD subunits. The previously enigmatic GATAD2 controls the asymmetry of the complex and directly recruits the ATP-dependent CHD remodeler. Unexpectedly, the MTA-MBD interaction acts as a point of functional switching. The transcriptional regulator PWWP2A modulates NuRD assembly by competing directly with MBD for binding to the MTA-HDAC-RBBP subcomplex, forming a ‘moonlighting’ PWWP2A-MTA-HDAC-RBBP complex that likely directs deacetylase activity to PWWP2A target sites. Taken together, our data describe the overall architecture of the intact NuRD complex and reveal aspects of its structural dynamics and functional plasticity.

## INTRODUCTION

The physical organization of DNA is a critical determinant of genome function. ATP-dependent chromatin remodeling enzymes use a conserved DNA translocase domain to alter the positions, occupancy and composition of nucleosomes, thereby regulating the availability of DNA for transcription, replication or repair. Chromatin remodelers typically exist as large multi-subunit complexes *in vivo*, and despite recent high-resolution structures of nucleosomes bound to the INO80 (1, 2) and SWR1 complexes (3), as well as to the Snf2 (4) and Chromodomain-Helicase-DNA-binding 1 (CHD1) remodelers (5), our understanding of how such enzymes bring about remodeling is still underdeveloped. This is particularly true for the nucleosome remodeling and deacetylase (NuRD) complex.

The NuRD complex is widely distributed among Metazoans and is expressed in most, if not all, tissues. It is essential for normal development (6, 7) and is a key regulator in the reprogramming of differentiated cells into pluripotent stem cells (8–10). Age-related reductions in NuRD subunit levels are strongly associated with memory loss, metastatic potential in human cancers (11), and the accumulation of chromatin defects (12, 13).

The mammalian NuRD complex comprises at least six subunits (**Figure S1a**), and for each subunit there are at least two paralogues, giving the potential for significant compositional heterogeneity. CHD4 (and its paralogues CHD3 and −5) is the ATP-dependent DNA translocase in the complex and harbours several regulatory and targeting domains. For example, the PHD domains of CHD4 can recognize histone H3 N-terminal tails bearing methyllysine marks (14–16), and the HMG domain has been shown to bind to poly-ADP(ribose) (17). What distinguishes NuRD from many other remodelers is that it harbours a second catalytic activity, imparted by the histone deacetylases HDAC1 and −2. MBD2 and −3 can bind hydroxymethylated and/or methylated DNA (18–20), and RBBP4 and −7 can each bind histone tails (21) and other transcriptional regulators (22, 23). The metastasis-associated proteins MTA1, −2 and −3 contain several domains that are associated with nucleosome recognition, whereas GATAD2A and GATAD2B bind to both MBD2 and −3 (24, 25) and CHD proteins (25, 26) but otherwise do not have known functions.

Some structural information is available for portions of the NuRD complex (**Figure S1b**): (i) HDAC1 forms a 2:2 complex with an N-terminal segment of MTA1 (MTA1_162–335_ – the ELM-SANT region) (27); (ii) single particle electron microscopy (SPEM) and X-ray crystallography data show that two copies of RBBP4 can bind the C-terminal portion of MTA1 (28–30); (iii) MBD2 and GATAD2A form a heterodimeric coiled-coil (24, 31); and (iv) a cryo-EM structure has been determined for the complex between the catalytic domain of CHD4 and a nucleosome particle (32). However, no structural data at any resolution have been presented for the intact complex. The subunit stoichiometry is also uncertain; recent studies using label-free mass spectrometry (33–38) have yielded variable results that are often at odds with the stoichiometries demonstrated in the known subcomplex structures.

The mechanisms by which NuRD selects target sites are also poorly understood. Transcriptional regulators such as FOG1 (22, 39) and BCL11A (40, 41) can bind to NuRD via the RBBP subunits but this mechanism is likely to account for only a small proportion of NuRD-genome interactions. Recently, we demonstrated that the chromatin-binding protein PWWP2A, which can selectively recognize H2A.Z-containing nucleosomes (42, 43), interacts robustly with the MTA, HDAC and RBBP subunits of NuRD. Surprisingly, however, GATAD2, MBD and CHD are not detected as interaction partners in these affinity-purification based experiments, suggesting that the two enzymatic activities in NuRD might in some situations be separable.

Against this background, we have used structural, biophysical and biochemical data to define the architecture of the NuRD complex. We first determine the subunit stoichiometry of the complex, using external peptide standards to provide a substantial increase in accuracy over previous measurements. Our findings corroborate all existing structural data for NuRD subcomplexes, pointing to a 4:2:2:1:1:1 ratio of RBBP, HDAC, MTA, MBD, GATAD2 and CHD subunits. We then demonstrate that the complex displays considerable conformational dynamics and we identify stable subcomplexes within NuRD, showing that the full complex is composed of two parts with separable enzymatic activities. A symmetric module comprising MTA, HDAC and RBBP subunits (MTA-HDAC-RBBP) carries the lysine deacetylase activity, whereas the MBD, GATAD2 and CHD subunits (MBD-GATAD2-CHD) act together to introduce DNA translocase activity into the complex. Our data indicate that, despite the underlying dimeric structure of the deacetylase module, the presence of the CHD-containing translocase module confers overall asymmetry in the complex. The connection between the modules is coordinated by the GATAD2 subunit and, unexpectedly, the interface between MTA-HDAC-RBBP and MBD-GATAD2-CHD is a site of regulation by additional proteins. In this context, we show that the co-regulator PWWP2A is able to compete directly with the translocase module for binding to the MTA-HDAC-RBBP subcomplex.

## RESULTS

### Subunit stoichiometry measurements partition the NuRD complex into symmetric and asymmetric modules with distinct catalytic activities

We purified native mammalian NuRD complex from murine erythroleukemia cells using our previously established protocol (**Figure 1a**, (44)). This strategy also yields a complex lacking the CHD subunit, which we have previously termed the Nucleosome Deacetylase (NuDe) complex. To interrogate the subunit composition of these complexes, we carried out data-independent acquisition mass spectrometry (DIA-MS) (45) using subunit-specific ^13^C/^15^N-labelled peptides as internal standards. This approach provides rigorous quantification of NuRD subunits together with information on paralogue composition (**Supplementary Results, Figure S1c**).

**Figure 1.**
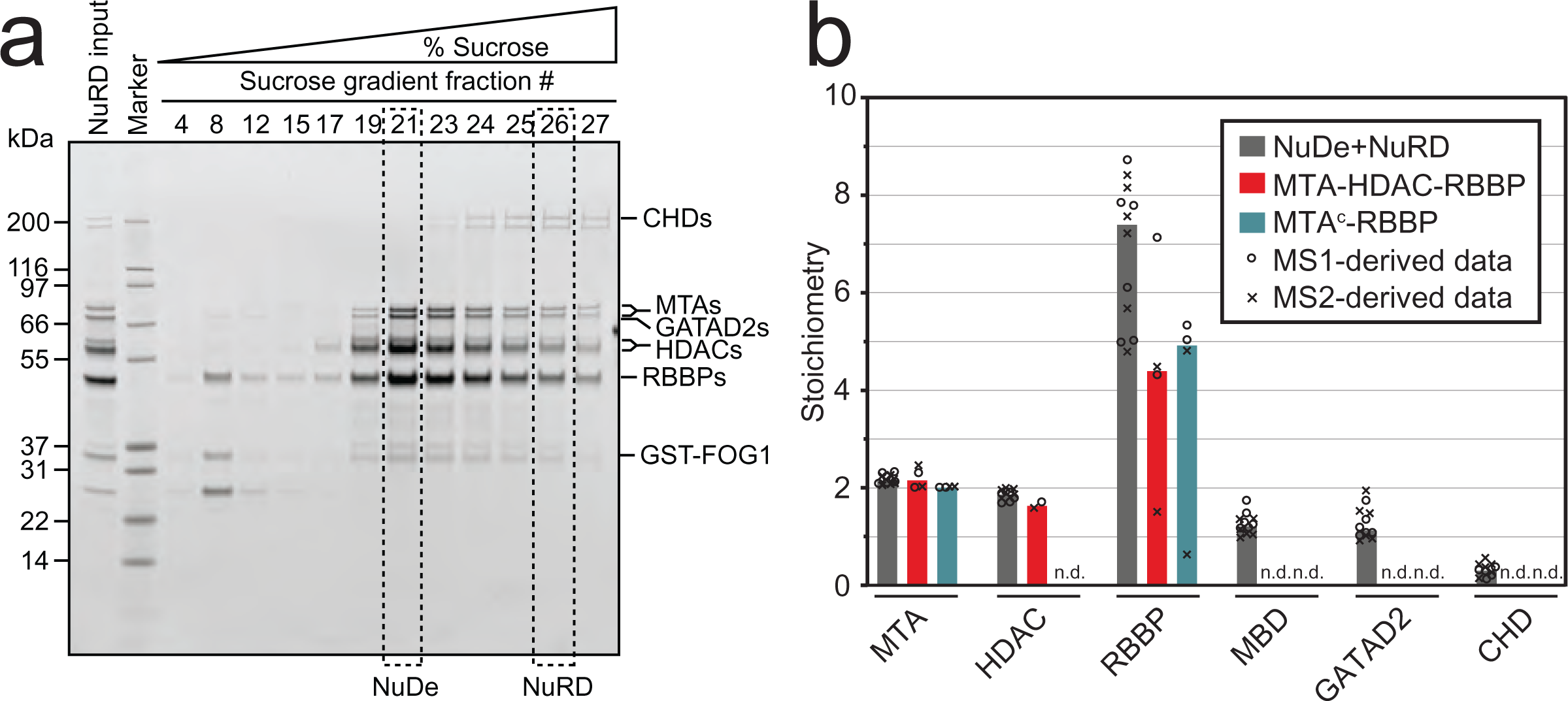
Absolute quantification of NuRD subunit stoichiometry by mass spectrometry. **a.** SDS-PAGE showing the NuRD complex after FOG1 pulldown (*NuRD input*), and showing fractions enriched for NuDe and NuRD following subsequent sucrose-gradient centrifugation. MBD proteins do not stain as well as other subunits. **b.** NuDe + NuRD, MTA-HDAC-RBBP and MTA^C^-RBBP subunit stoichiometry derived from DIA-MS data. Numbers are shown relative to the MTA/HDAC average, which is set to 2 of each based on the MTA1_162–335_-HDAC1 crystal structure (27). Open circles and crosses indicate the individual data points, from each biological replicate, derived from MS1 and MS2, respectively. The bars show the median value. “n.d.” indicates a null value.

For both NuDe and NuRD, we derived ratios of ∼2:2:7:1:1 for MTA:HDAC:RBBP:MBD:GATAD2 (**Figure 1b**, **Supplementary Data 2**). CHD4 was consistently sub-stoichiometric, reflecting the fact that complete separation of NuDe and NuRD was not possible because of overlap in their sedimentation profiles. The numbers are calculated relative to an MTA:HDAC ratio of 2:2 (**Supplementary Results**), based on the crystal structure of this complex (27).

Of all subunits, RBBP displayed the highest variability between samples (**Figure 1b**). The ratio of 7 RBBPs per NuRD complex was unexpected; however, it is the FOG1-RBBP interaction that was used as the affinity purification ‘handle’, most likely leading to an excess of NuRD-free RBBPs. Given this variability, we sought to more rigorously define the RBBP content of the complex. Previous structural and biochemical work shows that it is the C-terminal half of MTA1 that is responsible for recruiting RBBPs to the NuRD complex (25, 28, 30). To corroborate this conclusion, we expressed and purified a subcomplex comprising HDAC1, the N-terminal half of MTA2 (residues 1–429; MTA^N^) and MBD3_GATAD2CC_ (MTA^N^-HDAC-MBD_GATAD2CC_, **Figure S2a**). MBD3_GATAD2CC_ is full-length MBD3 stabilized by fusion to the coiled-coil domain of murine GATAD2A (residues 133–174), with which it dimerizes (46) (**Figure S1a**). Quantitative DIA-MS analysis showed that this MTA^N^-HDAC-MBD_GATAD2CC_ complex contains little or no RBBP protein, consistent with the idea that RBBP subunits are recruited to the complex solely by the C-terminal half of MTA (**Supplementary Data 2**). We confirmed this conclusion by expressing an MTA^C^-RBBP subcomplex comprising RBBP4 and the C-terminal half of MTA1 (residues 449–715) (**Figure S2a**); DIA-MS analysis showed that this complex has a stoichiometry of ∼1:2.5 (**Figure 1b**, **Supplementary Data 2**) – much closer to the expected 1:2 ratio. We likewise co-expressed and purified a subcomplex comprising MTA2, HDAC1 and RBBP7 (MTA-HDAC-RBBP, **Figure S2a**), which yielded a subunit ratio of ∼2:2:4 (**Figure 1b**, **Supplementary Data 2**).

As an orthogonal approach, we directly measured the molecular mass of the NuDe and NuRD complexes using size exclusion chromatography coupled to multiangle laser light scattering (SEC-MALLS). In both cases, we observed masses that were within 7% of the expected mass for a 2:2:4:1:1:1 complex (MTA:HDAC:RBBP:MBD:GATAD2:CHD4, **Figure S3a, Supplementary Results)**. Taken together, our data argue that the mammalian NuRD complex has a stoichiometry of 2:2:4:1:1:1 (MTA:HDAC:RBBP:MBD:GATAD2:CHD4). By integrating these findings with published biophysical and structural work on Drosophila NuRD and several subcomplexes (24, 27–30, 47), we can confidently conclude that NuRD is built from a symmetric 2:2:4 MTA-HDAC-RBBP deacetylase module and an asymmetric 1:1:1 MBD-GATAD2-CHD remodelling module.

The DIA-MS data also allowed us to quantify paralogue abundance in the native NuRD and NuDe complexes isolated from MEL/HEK293 cells. As shown **Figure S1c**, a distinct preference for HDAC1 over HDAC2 is observed, whereas MTA3 or MBD2 are nearly absent from our complex. The two GATAD2 paralogues are similarly abundant. Little or no CHD3 or CHD5 were detected in our samples.

### XLMS data establish the core architecture of the NuRD complex

We next asked how the deacetylase and remodeling modules of NuRD are physically connected. Our prior work shows that the only interaction linking these two modules is between MTA and MBD; no direct interactions between the deacetylase module (MTA-HDAC-RBBP) and either GATAD2 or CHD were identified (25). Our ability to co-express and purify a stable MTA^N^-HDAC-MBD_GATAD2CC_ subcomplex (see above) also points towards MBD bridging the two halves of the complex, and shows that neither the C-terminal half of MTA nor the RBBPs are required to couple the two halves of the complex. A structure of the MBD-GATAD2 complex (24) indicates that these two proteins directly interact, meaning that GATAD2 is recruited to the complex by MBD. Consistent with this idea, co-expression of MTA2, HDAC1, MBD3 and GATAD2A yields a stable complex containing all four proteins (MTA-HDAC-MBD-GATAD2, **Figure S2a,b**). Notably, this complex recruits significant amounts of endogenous CHD, whereas the MTA^N^-HDAC-MBD_GATAD2CC_ complex does not (**Figure S2a,b**). We therefore conclude that the architecture of NuRD consists of a dimeric deacetylase core of MTA, HDAC and RBBP subunits that directly interacts with MBD; in turn MBD binds GATAD2 and it is GATAD2 alone that dictates the recruitment of CHD and ultimately confers DNA translocase activity on the full complex (**Figure 2a**).

**Figure 2.**
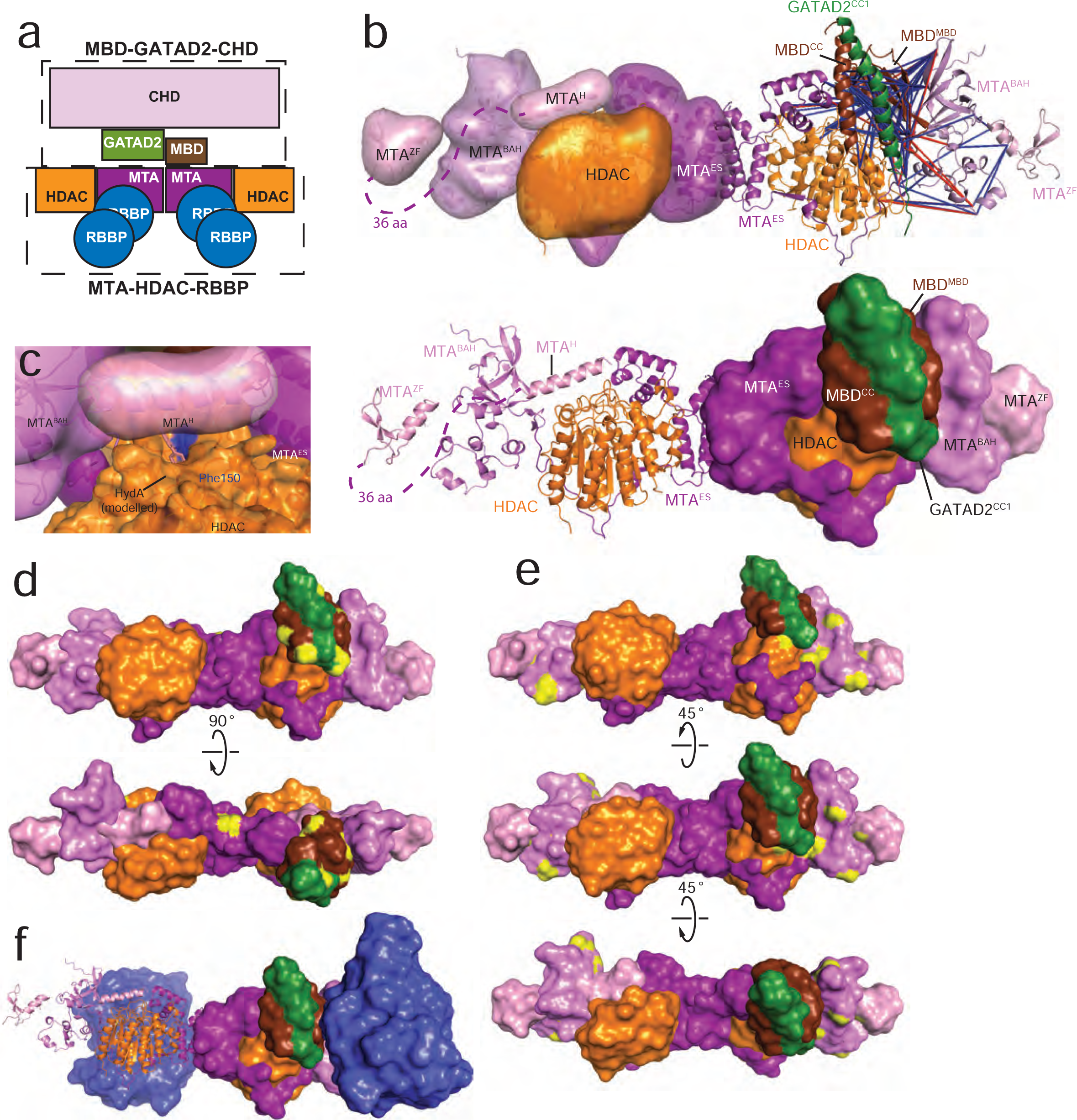
XLMS data for NuRD and NuRD subcomplexes. **a.** Schematic representation of NuRD. The two enzymatic modules (MTA-HDAC-RBBP and MBD-GATAD2-CHD) are indicated by dashed boxes. The MBD subunit bridges the two modules of the NuRD complex. **b.** Two representations of a molecular model derived from XL-driven rigid-body docking in HADDOCK. Domains used in the model include the MTA1_164–333_-HDAC dimer of heterodimers (MTA^ES^-HDAC), the MBD domain of MBD3_1–68_ (MBD^MBD^), a model of the MTA1_1–163_ BAH domain (MTA1^BAH^), a model of the MTA1_389–431_ GATA-type zinc-finger domain (MTA^ZF^), a model of the predicted MTA1_334–352_ helix (MTA^H^), and the heterodimeric coiled-coil formed by MBD3_216–249_ (MBD^CC^) and GATAD2A_136–178_ (GATAD2^CC1^). XLs satisfied (*blue*) and not satisfied (*red*) in this model are shown. **c.** Relative positions of the Phe150 (*blue*) in the active site of HDAC1, the MTA1_334–352_ helix (MTA^H^), and a modelled hydroxamic acid (HydA) HDAC inhibitor. The tight space formed between the MTA^H^ and HDAC1 suggests that MTA1 may restrict and modulate access to the HDAC active site. **d.** Residues forming XLs between the MBD3_69–216_ intrinsically disordered region (MBD^IDR^) and the MTA^N^-HDAC-MBD-GATAD2 core. The positions of the crosslinked residues on the core complex (*yellow*) point to the likely position of the MBD^IDR^ region of MBD3. **e.** Residues forming XLs between the MTA^C^-RBBP_2_ and the MTA^N^-HDAC-MBD-GATAD2 core. The positions of the crosslinked residues on core complex (*yellow*) provide clues on the approximate position of the MTA^C^-RBBP_2_ unit. **f.** An XL-based model showing the likely position of the two MTA^C^-RBBP_2_ units relative to the MTA^N^-HDAC-MBD-GATAD2 core. The model was created by manually positioning the two MTA^C^-RBBP_2_ units.

To examine the placement of subunits and, in particular, how the MBD subunit couples the two halves of the complex, we carried out covalent crosslinking combined with mass spectrometry (XLMS) on NuRD and several subcomplexes. These experiments yielded 752 unique XLs that were highly conserved across MTA^N^-HDAC-MBD_GATAD2CC_, MTA-HDAC-MBD-GATAD2, MTA-HDAC-RBBP, NuDe and NuRD complexes (**Figure S4a, Supplementary Data 1**).We observed a high density of XLs between: (i) RBBP and the C-terminal half of MTA; (ii) HDAC and the N-terminal half of MTA; (iii) MBD and the N-terminal half of MTA; and (iv) the first coil-coiled region (∼residues 136–178) of GATAD2 and MBD3. All these pairs have been previously reported to be the direct points of contact in the complex (24, 25, 27–31).

Of these XLs, 174 are between pairs of residues in known structures of subunits and subcomplexes. Ninety-four percent (range 88–100%) of these are consistent with the structural data (**Figure S4b, Supplementary Data 1**), providing a strong argument that these subcomplex structures recapitulate the architecture of the full NuRD complex in solution.

A set of 120 XLs connects pairs of domains that have known or readily modeled structures but for which relative positions in the NuRD complex are unknown. We have defined these domains as shown in **Figure S1**: MTA1_1–163_ (MTA1^BAH^: the BAH domain of MTA), the dimer of heterodimers HDAC1 + MTA1_164-333_ (MTA^ES^: the ELM-SANT domains of MTA), MTA1_334–352_ (MTA^H^: a 20-residue predicted α-helix to which 19 XLs were observed), MTA1_389–431_ (MTA^ZF^: a predicted GATA-type zinc-finger domain), MBD3_1–68_ (MBD^MBD^: the methyl-DNA binding domain) and the heterodimeric coiled-coil formed by MBD3_216–249_ (MBD^CC^: the MBD3 coiled-coil domain) and GATAD2A_136–178_ (GATAD2^CC1^: the first coiled-coil domain of GATAD2A). XL-driven rigid-body docking in HADDOCK (48) yields a domain architecture that is consistent with 93% (112/120) of the XLs (**Figure 2b**). This model places the BAH domain at the distal ends of the MTA^ES^-HDAC dimer of dimers where it is juxtaposed with the MBD^MBD^ domain. In turn, the MBD-GATAD2 coiled-coil domain packs against the MBD^MBD^ domain, despite the two domains being separated by ∼145 residues of sequence that is disordered in isolation (46). Only one copy of MBD3 is placed in the model because of the complex stoichiometry dictated by our DIA-MS data.

In our model, the MTA^H^ helix lies directly adjacent to the HDAC active site. **Figure 2c** highlights the well-described phenylalanine at the entry to the active site (Phe150, *blue*) and we have indicated the degree of access to the active site by modelling in a known HDAC inhibitor, hydroxamic acid (based on a structure of HDAC8; PDB: 5FCW) (49). This observation suggests that the MTA subunits in NuRD could potentially modulate HDAC activity.

The MBD subunit harbours a region of ∼145 residues (residues 69-216, MBD^IDR^) that has been demonstrated to be disordered in isolation (46) but which is predicted to be ordered by programs such as PONDR (50). We observed 162 XLs to residues in MBD^IDR^, but only in the N-terminal half of the sequence (**Figure S4c**). Approximately half of the XLs were to other unstructured residues in MBD3 but 92 were to structured portions of HDAC, MTA^N^, GATAD2^CC1^, MBD^CC^ and MBD^MBD^, forming a relatively contiguous surface against which this region might pack (**Figure 2d**). Concordant with these data, Williams and colleagues demonstrated that the N-terminal two-thirds of this region (but not the C-terminal third) can immunoprecipitate HDAC2, MTA2 and RBBP4 and that point mutations in this region abrogate the interaction (46).

One hundred and seventy-seven XLs were observed within and between RBBP proteins and the C-terminal half of MTA (which encompasses the two RBBP binding sites R1 and R2 as well as a ∼100-residue region that is predicted to be disordered, **Figure S1a**). Of these, 69 XLs occur within previously published crystal structures of the MTA1^R1^-RBBP4 and MTA1^R2^-RBBP4 subcomplexes and were highly consistent with the structural data (63/69 or 91%, **Figure S4c**). An additional 7 XLs were between structured regions but not in published crystal structures and we were able to use HADDOCK, together with published structures, to generate a model of MTA^C^-RBBP_2_. However, while this model fitted within our published low-resolution electron microscopy (EM) map of the complex (28), only 4 out 7 XLs (∼57%) could be satisfactorily mapped within the crosslinker distance constraints (**Figure S4d**); this fulfilment rate is far lower than the 93% observed for the MTA^N^-HDAC-MBD-GATAD2 core complex model above and might reflect substantial dynamics in this region of the complex.

Finally, a further 60 XLs were observed between the C-terminal half of MTA and our HADDOCK-modelled core complex. Again, the XLs highlight a contiguous surface (**Figure 2e**, *yellow*) that indicates the likely location of the MTA^C^-RBBP_2_ subcomplex. These XLs allow us to represent the general position taken up by the two MTA^C^-RBBP_2_ units in a full MBD-HDAC_2_-MTA_2_-RBBP_4_ complex (**Figure 2f**). The remaining portion of the complex, namely CHD and the C-terminal half of GATAD2, displayed very few inter-subunit XLs, meaning that their locations cannot be confidently modelled. This lack of XLs suggests that the CHD-GATAD2 subcomplex also displays considerable dynamics.

### GATAD2 controls the asymmetry of the NuRD complex

Our model indicates that there are two equivalent sites on the MTA-HDAC subcomplex that could accommodate an MBD subunit, raising the question as to why only one MBD is observed in the full NuRD complex. We therefore made DIA-MS measurements on the MTA^N^-HDAC-MBD_GATAD2CC_ subcomplex. Unexpectedly, we derived a 2:2:2 stoichiometry (**Figure 3a**) – consistent with the symmetry of the HDAC-MTA-RBBP subcomplex and our crosslink-directed modelling but inconsistent with the native stoichiometry we determined for NuRD. The finding that *in vitro* reconstituted MTA^N^-HDAC-MBD_GATAD2CC_ forms a 2:2:2 complex was corroborated by SEC-MALLS data (**Figure 3b**), which yielded a MW of ∼290 kDa, in close agreement with the 288-kDa mass predicted. FLAG-tagged MBD3 can also immunoprecipitate untagged MBD3 when both are co-expressed with HDAC and MTA, confirming that there are two MBD-compatible binding sites within the one complex (**Figure S5**).

**Figure 3.**
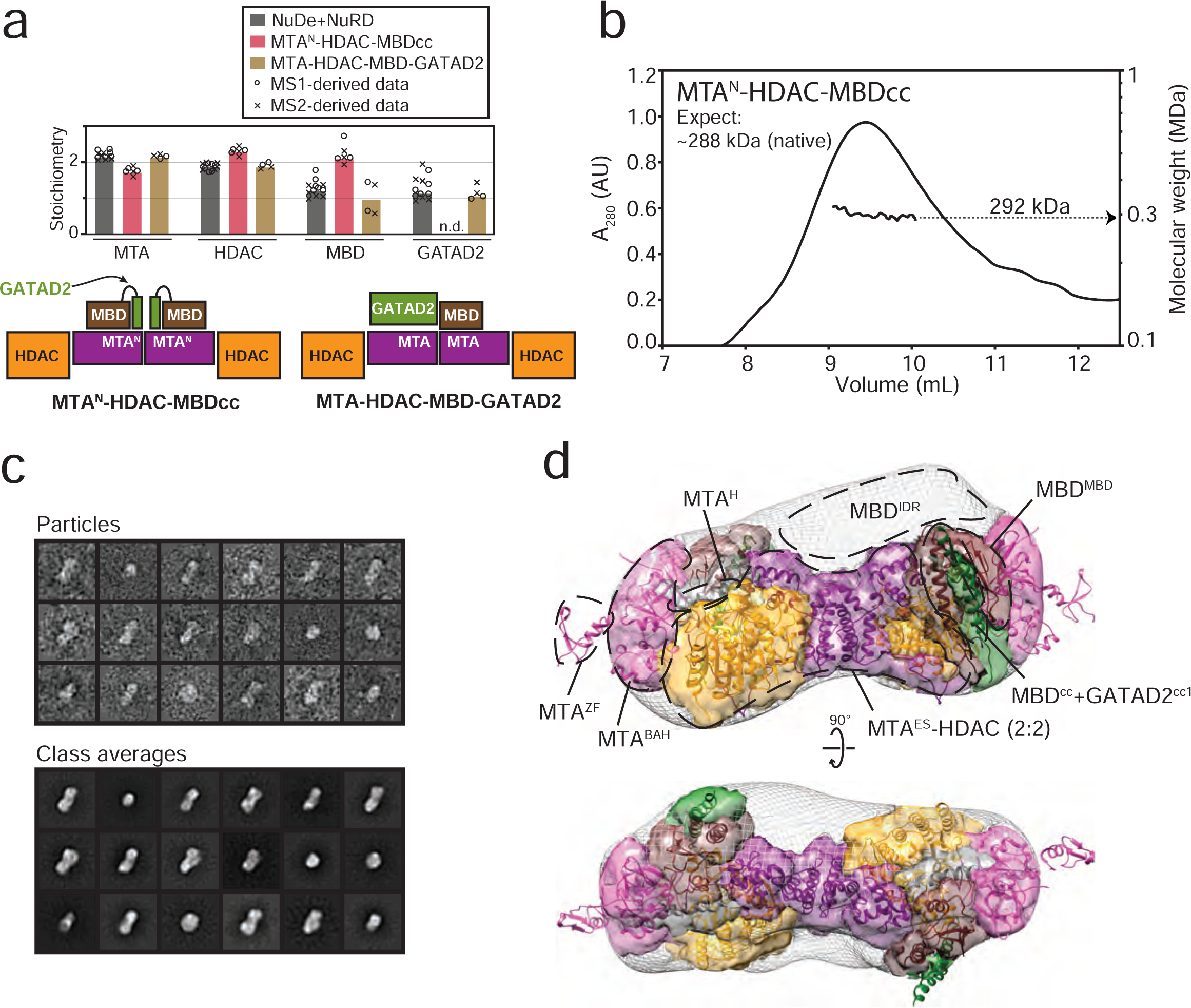
Structure of the MTA^N^-HDAC-MBD_GATAD2CC_ complex. **a.** *Top.* Subunit stoichiometry derived from DIA-MS data for MTA^N^-HDAC-MBD_GATAD2CC_ and MTA-HDAC-MBD-GATAD2 complexes. Data for NuDe+NuRD from Figure 1b are shown for comparison. Numbers are shown relative to the MTA/HDAC average, which is set to 2 of each based on the MTA1_162–335_-HDAC1 crystal structure (27). Open circles and crosses indicate the individual data points, from each biological replicate, derived from MS1 and MS2, respectively. The bars show the median value. “n.d.” indicates a null value. *Bottom*. Schematics of MTA^N^-HDAC-MBD_GATAD2CC_ and MTA-HDAC-MBD-GATAD2 for reference. **b.** SEC-MALLS experiment for MTA^N^-HDAC-MBD_GATAD2CC_ under non-denaturing conditions. **c.** Selected reference-free 2D class averages for MTA^N^-HDAC-MBD_GATAD2CC_. **d.** Final 3D envelope for MTA^N^-HDAC-MBD_GATAD2CC_ refined to 29 Å resolution. Also shown is the molecular model of the complex adapted from Figure 2b to include a second molecule of MBD_GATAD2CC_ with two-fold symmetry. Subunit colours are as in Figure 2. The wire mesh indicates regions of the map for which no model density is currently assigned; these regions could accommodate the C-terminal ∼100 residues HDAC and ∼145 residues of MBD3^IDR^, for which no structure is available.

Using negative stain electron microscopy (EM), we recorded 325 micrographs of the purified MTA^N^-HDAC-MBD_GATAD2CC_ complex (**Figure 3c and S2c**). From these micrographs we generated class averages that represent different orientations of the complex. The dataset was relatively homogeneous, so using 3D classification and refinement routines in RELION, we obtained a low-resolution structural envelope (29 Å according to the FSC 0.143 criterion) that reveals a bi-lobed structure with an approximate two-fold symmetry (**Figure 3d**), in line with our stoichiometry data. The model derived from our XLMS data is overall well accommodated within this density envelope (**Figure 3d**). Only the MTA^ZF^ domain significantly protrudes from the map, perhaps indicating that the ZF domain is dynamic; this is likely, given it is connected to the rest of MTA^N^ by a 36-residue linker that is predicted to be disordered. The envelope presented in **Figure 3d** has been segmented and colored according to occupancy by the various structural domains that are represented in our XLMS-derived model. The wire mesh indicates regions of the map for which no model density is currently assigned. These are likely indicative of regions of sequence for which no structures are available, specifically the C-terminal ∼100 residues of HDAC and ∼145 residues of MBD^IDR^. Encouragingly, the largest part of this unassigned density lies in the position predicted for MBD^IDR^ from our XLMS data (**Figure 2d and 3d**).

The unexpected 2:2:2 stoichiometry of the MTA^N^-HDAC-MBD_GATAD2CC_ complex raised the question of why only one MBD is found in the intact NuDe/NuRD complexes. We hypothesized that it is the presence of the GATAD2 subunit that prevents binding of a second MBD subunit in the larger complexes. Indeed, when we co-expressed GATAD2A with MTA2, HDAC1 and MBD3 (the MTA-HDAC-MBD-GATAD2 complex, **Figure S2a**), the expected 2:2:1:1 stoichiometry was restored, as judged by DIA-MS (**Figure 3a**, **Supplementary Data 2**). We therefore conclude that the GATAD2 subunit is responsible not only for recruiting CHD remodelling activity to the NuRD complex but also for controlling the asymmetry of the complex.

### RBBP subunits introduce substantial conformational dynamics into the NuRD, NuDe and MTA-HDAC-RBBP complexes

We next analysed three other (sub)complexes by negative-stain EM. Datasets were collected for the MTA-HDAC-RBBP, NuDe and NuRD complexes (**Figure S2c** and **S6**). In each case, particles were observed that could be aligned and classified to produce 2D class averages (**Figure 4a**). Examination of the classes obtained for MTA-HDAC-RBBP made it clear that significant shape heterogeneity exists, even for particles of a similar apparent size. Given the relative homogeneity of the MTA^N^-HDAC-MBD_GATAD2CC_ complex preparation, we conclude that the MTA^C^-RBBP_2_ units exhibit considerable dynamics relative to the MTA^N^-HDAC-MBD_GATAD2CC_ core. Because of this dynamic behavior, we did not generate 3D maps but rather restricted our analysis to the 2D classes, which we interpreted as projections of conformational variants of the complex. **Figure 4b** shows that 2D classes for the MTA-HDAC-RBBP complex can be readily interpreted as comprising a central MTA^N^-HDAC module (magenta and orange, EMD-3399, (30)) and two MTA^C^-RBBP_2_ modules (blue, EMD-3431, (28)) that can rotate relative to the MTA^N^-HDAC core. The only difference between the models shown is the relative rotation of the MTA^C^-RBBP_2_ units.

**Figure 4.**
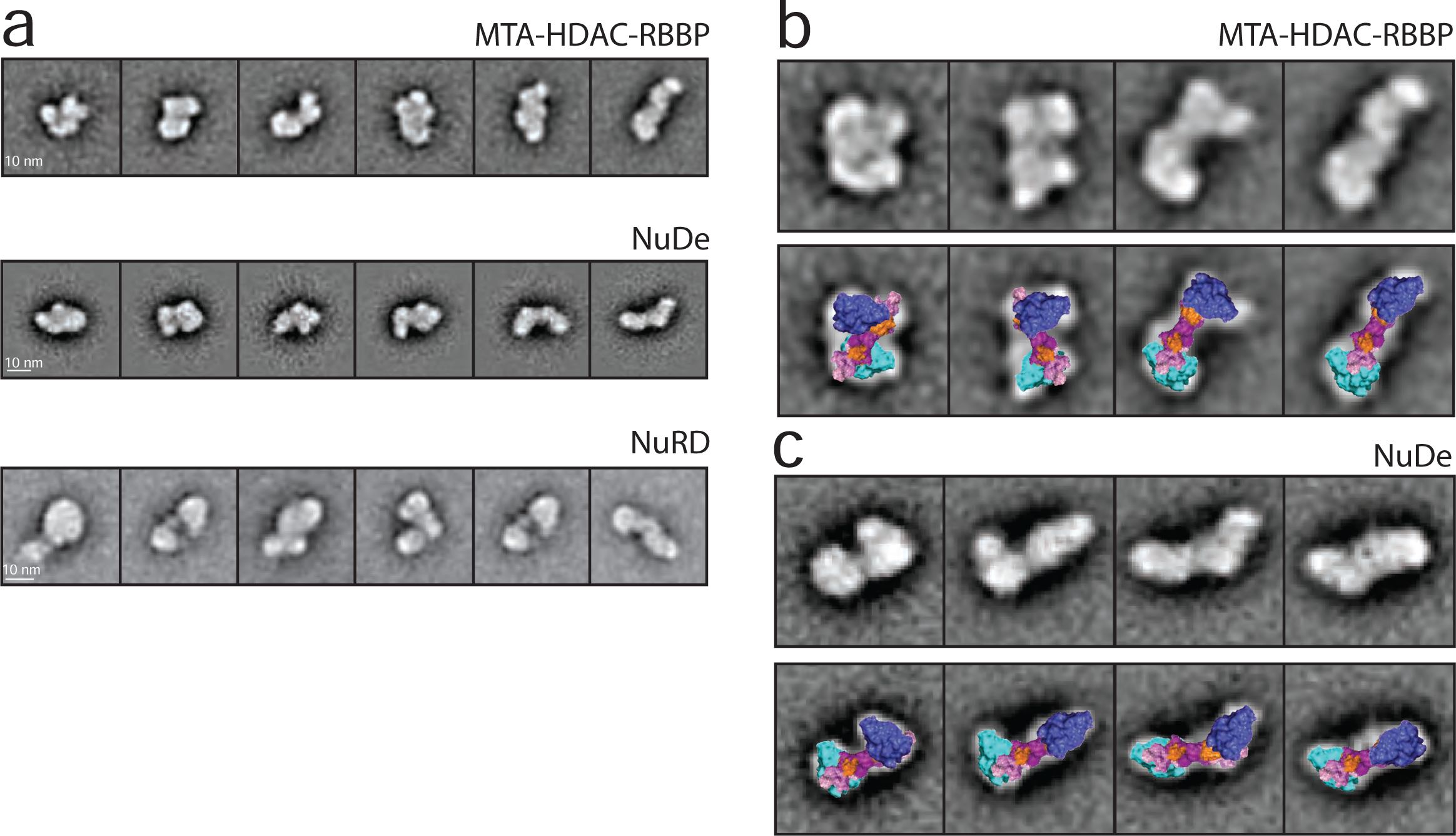
RBBP-containing complexes display significant shape heterogeneity. **a.** 2D classes for MTA-HDAC-RBBP, NuDe and NuRD complexes. **b, c.** Significant dynamics are observed for the MTA-HDAC-RBBP and NuDe complexes. *Upper panels.* Selected 2D classes for the indicated complexes. *Lower panels.* The same classes, but overlaid with models that comprise the MTA^N^-HDAC core from Figure 2b (*magenta* and *orange*, EMD-3399, (30)) and two MTA^C^-RBBP_2_ modules (blue, EMD-3431, (28)) that can rotate relative to the MTA^N^-HDAC core (magenta and orange, EMD-3399, (30)).

Analysis of the NuDe data revealed a similar pattern (**Figure 4c**). GATAD2 is likely to be highly dynamic and therefore does not give rise to clear additional density. Interpretation of the NuRD data is more challenging, as there is a lack of reliable 3D structural data for most of CHD4. Nonetheless, a similar type of conformational heterogeneity is apparent (**Figure 4a**), as expected for our model.

### PWWP2A competes directly with MBD-GATAD2-CHD for binding to the MTA-HDAC-RBBP module of NuRD

Our data demonstrate that MBD and GATAD2 form a nexus that connects two symmetry-mismatched modules of NuRD – namely the remodelling and the deacetylase modules. This finding raises the question as to whether either module has a physiological role outside of the NuRD complex. We and others recently demonstrated that PWWP2A, a transcriptional regulator that is important for neural crest differentiation, can immunoprecipitate HDAC, MTA and RBBP but not other NuRD subunits (42, 51). PWWP2A therefore appears to selectively bind MTA-HDAC-RBBP but not intact NuRD. To better understand this interaction, we expressed FLAG-PWWP2A with combinations of HDAC1, MTA1, MBD3 and GATAD2B and carried out pull-downs. As expected, FLAG-PWWP2A pulled down MTA1 and HDAC1 (**Figure 5a, *lane 5***). Strikingly, however, the additional co-expression of either MBD3 or a combination of MBD3 and GATAD2B (***lanes 4 and 6***, respectively) made no difference – only HDAC1 and MTA1 were retained by PWWP2A. In contrast, when PWWP2A was absent, MBD3 and GATA2B *did* form a complex with the MTA-HDAC core, as expected (**Figure 5b, *lanes 3 and 4***). These data demonstrate that PWWP2A competes directly with MBD for binding to the deacetylase module of NuRD.

**Figure 5.**
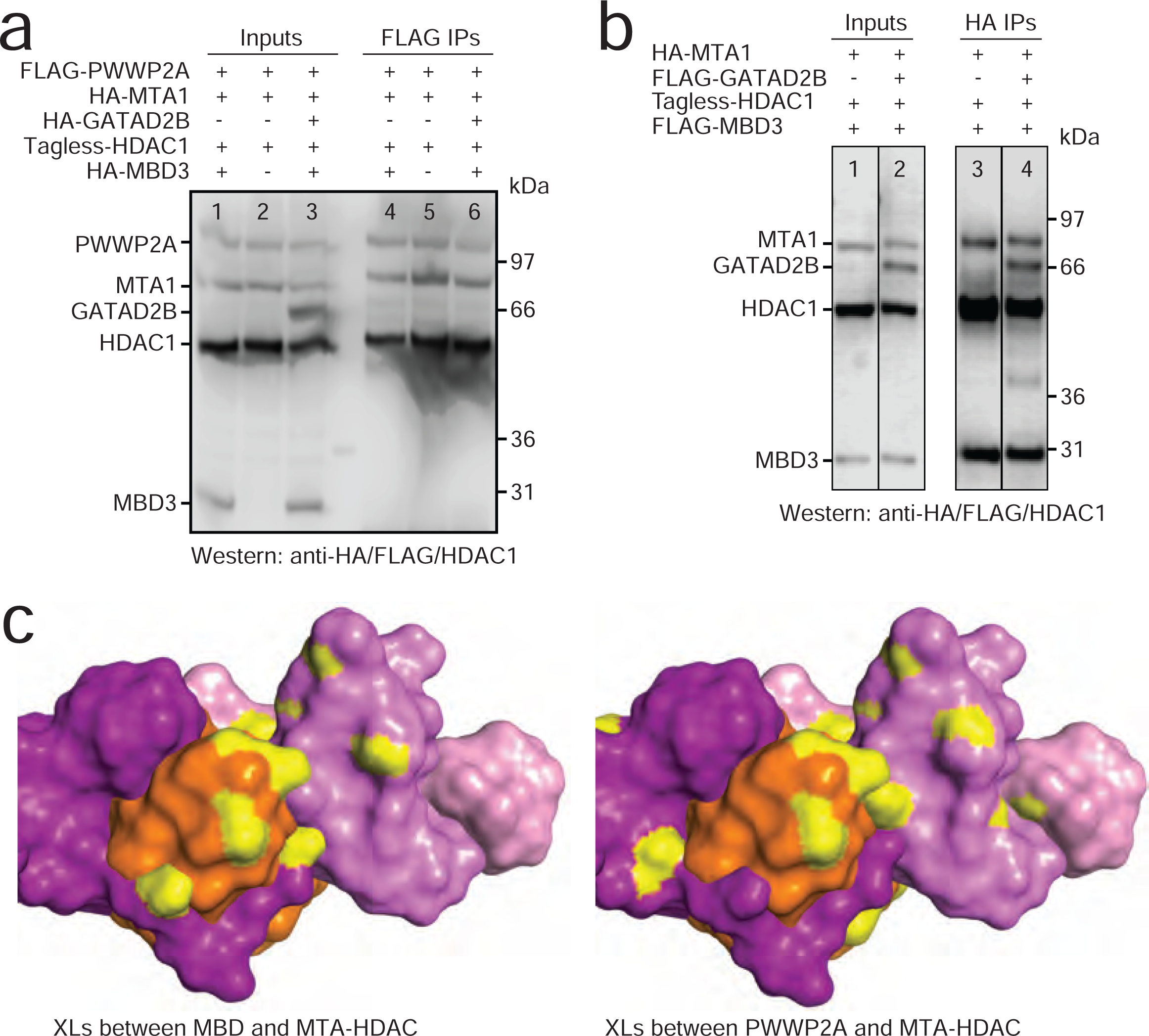
PWWP2A competes with MBD3/GATAD2B for binding to the MTA-HDAC-RBBP complex. **a.** Pulldowns showing that FLAG-PWWP2A purified on anti-FLAG beads pulls co-expressed HDAC1 and MTA1 out of cell lysate (*lanes 2 + 5*), but that neither co-expressed MBD3 (*lanes 1 + 4*) nor a combination of co-expressed MBD3 and GATAD2B (*lanes 3 + 6*) are pulled down. All NuRD components were HA tagged except for HDAC1, which was untagged. **b.** HA pulldown showing that co-expressed HA-MTA1, HDAC1, FLAG-MBD3 and FLAG-GATAD2B can form a single complex (MTA-HDAC-MBD-GATAD2). **c.** Comparison of residues in the MTA-HDAC complex that exhibit XLs to either MBD (*left*) or PWWP2A (*right*).. Colours: MTA (*magenta*), HDAC (*orange*) and crosslinked residues (*yellow*).

To corroborate this finding, we collected XLMS data on a PWWP2A-MTA-HDAC-RBBP complex (**Supplementary Data 1**). 146 unique XLs were observed between PWWP2A and either HDAC or the N-terminal half of MTA. **Figure 5c** shows that the set of residues in MTA1 and HDAC1 that crosslink to PWWP2A closely matches those that crosslink to MBD in the MTA^N^-HDAC-MBD_GATAD2CC_ complex. These data together support the conclusion that PWWP2A directly competes with MBD for binding to a common surface on the MTA-HDAC core complex. In an *in vivo* context, this provides a mechanism by which PWWP2A can recruit the deacetylase activity of NuRD to specific genomic loci without co-recruiting the remodelling activity of the CHD subunit.

## DISCUSSION

### NuRD assembly is regulated at an interface between separable histone deacetylase and DNA translocase units

The data presented here define the subunit stoichiometry of the mammalian NuRD complex, elucidate key features of NuRD architecture and demonstrate that the NuRD complex is built from two separable structural entities that have distinct catalytic activities. One of these entities – MTA-HDAC-RBBP – has two-fold symmetry, can be readily purified and harbours the histone deacetylase activity. In contrast, the other – MBD-GATAD2-CHD – has a 1:1:1 stoichiometry, is inherently asymmetric, and contributes the ATP-dependent remodeling activity.

A central question regarding the architecture of the NuRD complex has been how the symmetric 2:2:4 MTA-HDAC-RBBP module and the inherently asymmetric 1:1:1 MBD-GATAD2-CHD unit are connected. Our data show that MBD alone is insufficient to ‘break the symmetry’ of MTA-HDAC-RBBP and it is the recruitment of GATAD2 that leads to asymmetry in the full complex. GATAD2 is also responsible for the recruitment of a CHD subunit to the complex (25), via an interaction that is mediated by the GATA-type zinc finger of GATAD2 (26). We do not observe GATAD2 binding to MTA, HDAC or RBBP (25), and thus the means by which GATAD2 prevents a second copy of MBD from binding to MTA-HDAC-RBBP remains unclear. It is possible that the binding of GATAD2 to one MBD sterically blocks access to the second MTA subunit without involving a direct GATAD2-MTA interaction.

The modular nature of the NuRD complex is functionally relevant. We have shown recently that the coregulator PWWP2A can selectively bind the MTA1-HDAC-RBBP module and likely deacetylates H3K27 and H2A.Z (42). Here, we demonstrate that the PWWP2A-MTA1-HDAC-RBBP interaction effectively rejects the MBD-GATAD2-CHD subunits by competing directly with MBD for binding to a common surface on MTA. Recent work on PWWP2A/B suggests that two copies of PWWP2A/B interact with MTA-HDAC-RBBP (51), mirroring the 2:2:2 stoichiometry we see for MTA-HDAC-MBD. Thus, the MTA-HDAC-RBBP module appears to have a cellular function independent of the intact NuRD complex, with the MTA subunit acting as a regulatory control point for competitive interactions that can lead to NuRD-independent activities for NuRD subcomplexes.

Our stoichiometry and XLMS data, combined with published findings, allows us to present a structural model for the region of the NuRD complex comprising MTA^N^, HDAC, MBD^MBD^, and the MBD^CC^-GATAD2^CC1^ coiled-coil domain. The model, which is additionally consistent with our low-resolution electron microscopy data, significantly extends our understanding of NuRD architecture and delineates the region that acts as the point of connection between the deacetylase and remodelling modules, as well as other interacting partners.

### The NuRD complex is dynamic

Our EM data indicate that the NuRD complex does not exist in a single well-defined conformation. The MTA-HDAC-RBBP, NuDe and NuRD complexes appear to exhibit substantial conformational heterogeneity, indicating that the RBBP subunits are a major source of structural dynamics. Consistent with this conclusion, our quantitative MS data and published structural data (29, 30) also suggest that the two MTA^C^-RBBP_2_ modules can undergo significant movement relative to the MTA^N^-HDAC core. The model shown in **Figure 6a** indicates the likely extent of motion exhibited by the MTA^C^-RBBP_2_ units, based on the XLMS and EM data. This mobility could have a role in docking the NuRD complex (or the competing PWWP2A-MTA-HDAC-RBBP complex; **Figure 6b**) onto nucleosomes via histone H3 (30), in preparation for deacetylation and/or remodeling. RBBPs also work as adaptors for interactions with transcriptional regulators (*e.g.*, FOG1 (22), ZNF827 (52), PHF6 (53)), which could help guide the NuRD complex to specific genomic sites.

**Figure 6.**
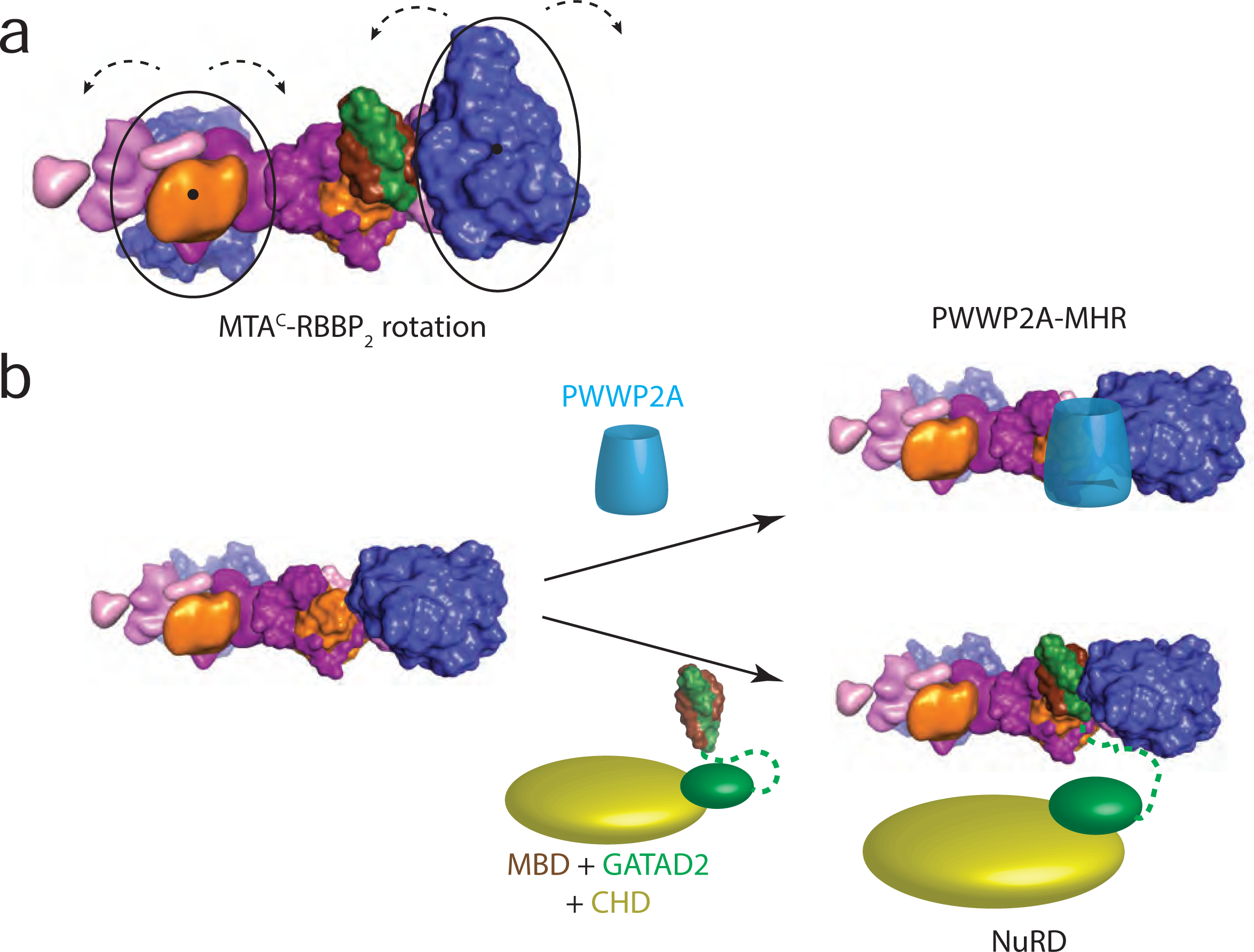
RBBP subunits introduce conformational dynamics and PWWP2A competes directly with the MBD-GATAD2-CHD module. **a.** A model based on XLMS and EM data depicting some of the possible range of movements by the MTA^C^-RBBP_2_ unit. **b.** PWWP2A competes directly with the MBD-GATAD2-CHD module for binding to the MTA-HDAC-RBBP module of NuRD, forming a PWWP2A-MTA-HDAC-RBBP complex. All NuRD subunit colours are as shown in Figure 2.

Considering that CHD and GATAD2 subunits together make up 25% of the mass of the NuRD complex, the lack of inter-subunit crosslinks to CHD and the C-terminal half of GATAD2 was unexpected and suggests that these subunits are even more dynamic than the RBBPs relative to the HDAC-MTA core. This conclusion is in accord with our EM data for the intact NuRD complex, which showed a smaller than expected increase in particle size compared to the subcomplexes. It is notable that the GATAD2 and CHD subunits have large regions of sequence that are predicted to be intrinsically disordered, and that the recent structure of CHD4 bound to a nucleosome (32) displayed density for only roughly a third of the protein.

The NuRD complex also displays considerable compositional dynamics and there are several examples of functional consequences of such dynamics. MBD2- and MBD3-specific NuRD complexes have distinct biochemical properties (Le Guezennec et al., 2006) and CHD3, −4 and −5 NuRD complexes operate at different stages of mammalian brain development (Nitarska et al., 2016). Furthermore, PWWP2A exhibits a preference for an MTA-HDAC-RBBP complex built on MTA1, rather than MTA2 (54). The NuRD complex that we isolate from MEL cells shows a distinct preference for HDAC1 over HDAC2 and both MTA3 and MBD2 are essentially absent from our complex (**Figure S1c**). The functional consequences of this subunit selectivity remain to be elucidated.

In summary, our data demonstrate that the NuRD complex is a highly dynamic assembly made from symmetric and asymmetric units that carry distinct and separable enzymatic activities. The interface between these units serves as a regulatory focal point in complex assembly, and direct competition with other transcriptional coregulators can lead to the MTA-HDAC-RBBP deacetylase module moonlighting independent of the full NuRD complex.

## METHODS

### Plasmids

Constructs used in this work have been previously described in (25, 28). Additional constructs for human GATAD2A (**<u>Q8CHY6</u>**) and human PWWP2A (**<u>Q96N64</u>**) were similarly cloned into the pcDNA3.1 expression vector to generate FLAG- and HA-tagged proteins.

### MTA-HDAC-RBBP, MTA^N^-HDAC-MBD_GATAD2CC_, MTA^C^-RBBP, MTA-HDAC-MBD-GATAD2, and PWWP2A-MTA-HDAC-RBBP expression and purification

Co-transfection, and transient overexpression of proteins in suspension HEK Expi293F^TM^ cells (Thermo Fisher), and purification by FLAG-affinity pulldowns were essentially performed as previously described in (25), with two exceptions: (i) that the cultures were scaled up to accommodate litre-sized cultures; and (ii) 3 mM ATP, 3 mM MgCl_2_ were added to the lysis and wash buffers. Selected elutions were pooled and concentrated using Amicon Ultra-2, 100 kDa MWCO devices (Merck Millipore), pre-blocked with 5% (v/v) Tween-20. Typical yields were 1–2 mg per litre of culture. The plasmid combinations used were FLAG-MTA2, tagless HDAC1 and HA-RBBP7 to express the MTA-HDAC-RBBP subcomplex, and FLAG-MBD3_GATAD2CC_, HA-MTA2^N^ and tagless HDAC1 for the MTA^N^-HDAC-MBD_GATAD2CC_ subcomplex, and FLAG-MTA2, HA-MBD3 and tagless HDAC1 and HA-GATAD2A for the MTA-HDAC-MBD-GATAD2 subcomplex, FLAG-MTA1^C^ and HA-RBBP4 for the MTA^C^-RBBP_2_ subcomplex, and FLAG-PWWP2A, HA-MTA1, tagless HDAC1 and HA-RBBP4 for the PWWP2A-MTA-HDAC-RBBP complex.

### NuDe and NuRD purification

Samples of NuDe and NuRD were isolated by FOG1 affinity pulldown from cultured mouse erythroleukemia (MEL) cells as described in (44).

### Sucrose density gradient ultracentrifugation and fractionation

Sucrose density gradients for both GraFix (55) and non-GraFix samples were performed essentially as described in (28). Either a 5.5 mL SW55 Ti or a 12 mL SW41 Ti Beckman Coulter rotor was used.

### Crosslinking-mass spectrometry (XLMS)

Sample preparation for XLMS using disuccinimidyl suberate (DSS) and adipic acid dihydrazide (ADH) crosslinkers were essentially as described previously (28). Crosslinking reactions using bissulfosuccinimidyl suberate (BS3) were performed essentially as described for DSS crosslinking with the exception that a 50 mM stock was made in Milli-Q water instead of dimethylformamide.

Post-sucrose gradient separation, fractions containing NuRD and NuDe complexes were pooled, buffer exchanged into sucrose-free buffers and concentrated using Vivaspin tricellulose acetate centrifugal filters (20 K MWCO; Sartorius). The centrifugal filters were pre-blocked with 0.1% (v/v) Tween-20. For each crosslinking experiment, ∼10–15 µg of the complex at a concentration of ∼0.1–0.15 mg/mL was used. For MTA-HDAC-RBBP, it is essentially the same as for NuRD and NuDe except that sucrose gradient separation was omitted.

For the MTA^N^-HDAC-MBD_GATAD2CC_, the MTA-HDAC-MBD-GATAD2 subcomplexes, and the PWWP2A-MTA-HDAC-RBBP complex, eluates were buffer exchanged into a buffer comprising 50 mM HEPES pH 7.5 and 150–250 mM NaCl using an Amicon Ultra 0.5 mL or 4 mL centrifugal filter (50 K MWCO; Merck Millipore). For each crosslinking experiment, ∼10–20 µg of subcomplex at a concentration of 0.2–1 mg/mL was used.

XLMS data for the MTA^C^-RBBP complex were recorded previously (28).

Sample preparation, LC-MS/MS data collection and database searches were essentially performed as described previously (28), with exceptions that mass analyses were performed using either a Q-Exactive or Q-Exactive Plus mass spectrometer (ThermoFisher Scientific). Mass spectrometer settings were essentially the same as previously published (28).

For the database searches, all taxonomies in the UniProt database (Nov 2013 – Jul 2016; 541,762– 551,705 entries) were searched.

### Analysis of XLMS data

Analysis of the XLMS data was performed with pLINK v2.3.5 (56). pLINK search parameters and data analysis were essentially the same as (28) with the following differences: Peptide mass between 600–10,000 Da and peptide length between 6–100 were considered, up to three missed cleavages were allowed, BS3/DSS crosslinking sites were Lys, Ser, Tyr, Thr and protein N-terminus. The default FDR of 5% was used. Only peptides with a precursor mass error of ≤ ±10 ppm, E-value scores of ≤1×10^−3^, and with at least four fragment ions on both the alpha- and beta-chain each were retained for further analysis.

For crosslinking schematics as shown in **Figure S4a**, the xVis webserver was used (57).

For modeling, the list of XLs was further filtered to remove redundant hits and crosslinked intra-protein residues of ≤10 residues apart were discarded. This final non-redundant list and unfiltered pLINK search outputs can be found in **Supplementary Data 1**. All mass spectrometry data and XLMS search results have been deposited to the ProteomeXchange Consortium via the PRIDE partner repository (58) with the dataset identifier PXD010111.

### Data-independent acquisition mass spectrometry and sample preparation

Proteotypic peptides (PTPs) selection is discussed in **Supplementary Results**. Selected peptides were synthesised with stable isotopes (carboxy-terminal Arg (^13^C_6_; ^15^N_4_) or Lys (^13^C_6_; ^15^N_2_)) as Absolutely Quantified SpikeTidesTM TQL with the cleavable Qtag by JPT Peptide Technologies. Peptides were supplied as 5 × 1 nmol aliquots and each 1 nmol aliquot per PTP were resuspended with 20% (v/v) acetonitrile in 80% (v/v) 0.1 M NH_4_HCO_3_, pooled, aliquoted, freeze-dried and stored at –20 °C. For each MS experiment, each aliquot was resuspended in 20% (v/v) acetonitrile in 80% (v/v) 0.1 M NH_4_HCO_3_, the required amount taken and the remaining excess peptide was then discarded.

Post-affinity chromatography and sucrose gradient fractionation, heavy-labelled synthetic peptides were added to the highly-purified NuRD or NuRD subcomplexes, and the samples were then freeze-dried. Once dried, the samples were then re-solubilised with 50 µL of 8 M urea dissolved in 50 mM Tris.Cl pH 8. The samples were then reduced (5 mM TCEP, 37 °C, 30 min) and alkylated (10 mM iodoacetamide, room temperature in the dark, 20 min). The samples were then diluted to 6 M urea with 50 mM Tris.Cl pH 8 and Trypsin/Lys-C mix (Promega) was added to a final enzyme:substrate ratio of 1:20 (w/w). The sample was then incubated for 37 °C for 4 h. Following this initial digestion step, the sample was further diluted to 0.75 M urea with 50 mM Tris.Cl pH 8 and additional Trypsin/Lys-C mix was added to a final enzyme:substrate ratio of 1:10 (w/w). The sample was then incubated at 37 °C overnight for ∼16 h. Following the overnight digestion, the samples were acidified with formic acid to a final concentration of 2% (v/v) and centrifuged (20,000 *g*, 10 min). The supernatant was then desalted using in-house made C18 stagetips.

All samples for stoichiometry analysis were performed using a Q-Exactive mass spectrometer. In every MS experiment/injection, 100 fmol of each reference heavy synthetic peptide was injected, unless otherwise stated. For every biological replicate that was analyzed, 1–2 replicates of DDA runs and 3–5 replicates of DIA runs were performed.

For LC-MS/MS, peptides were resuspended in 3% (v/v) acetonitrile, 0.1% (v/v) formic acid and loaded onto a 40–50 cm × 75 µm inner diameter column packed in-house with 1.9 µm C18AQ particles (Dr Maisch GmbH HPLC) using an Easy nLC-1000 UHPLC (Proxeon). Peptides were separated using a linear gradient of 5–30% buffer B over 120 min at 200 nL/min at 55 °C (buffer A consisted of 0.1% (v/v) formic acid; while buffer B was 80% (v/v) acetonitrile and 0.1% (v/v) formic acid). For DDA: after each full-scan MS1 (*R* = 70,000 at 200 *m/z*, 300–1750 *m/z*; 3 × 10^6^ AGC; 100 ms injection time), up to 20 most abundant precursor ions were selected for MS/MS (*R* = 35,000 at 200 *m/z*; 1 × 10^6^ AGC; 120 ms injection time; 30 normalized collision energy; 2 *m/z* isolation window; 8.3 × 10^5^ intensity threshold; minimum charge state of +2; dynamic exclusion of 60 s). For DIA: after each full-scan MS1 (*R* = 140,000 at 200 *m/z* (300–1600 *m/z*; 3 × 10^6^ AGC; 120 ms injection time), 16 × 25 *m/z* isolations windows in the 425–825 *m/z* range were sequentially isolated and subjected to MS/MS (*R* = 17,500 at 200 *m/z*, 3 × 10^6^ AGC; 60 ms injection time; 30 normalized collision energy). After the first eight 25 *m/z* isolations and MS/MS and before the last eight 25 *m/z* isolations, a full-scan MS1 was performed. 25 *m/z* isolation window placements were optimised in Skyline (59) to result in an inclusion list starting at 437.9490 *m/z* with increments of 25.0114 *m/z*.

### DIA data processing

All DDA data were processed using Proteome Discoverer v1.4 and searched with Sequest HT against all taxonomies in the UniProt database (Nov 2013 – Apr 2015; 541,762 – 548,208 entries). The data were searched with oxidation (M) and carbamidomethyl (C) as variable modifications using a precursor-ion and product-ion mass tolerance of ± 20 ppm and ± 0.02 Da. All results were filtered using Percolator to a false discovery rate of 1%.

All DIA data were processed using Skyline v3.5.0.9319 (59). Reference spectral libraries were built in Skyline with .msf files using the BiblioSpec algorithm (60). Precursor and product ion extracted ion chromatograms were generated using extraction windows that were two-fold the full-width at half maximum for both MS1 and MS2 filtering. Ion-match tolerance was set to 0.055 *m/z*. For MS1 filtering, the first three isotopic peaks with charges +2 to +4 were included while for MS2, *b-* and *y-* type fragments ions with charges +1 to +3 were considered. To ensure correct peak identification and assignment, the following criteria had to be met: (i) co-elution of light and heavy peptides; (ii) averaged dot product values of ≥0.95 between peptide precursor ion isotope distribution intensities and theoretical intensities (idotp) for both the heavy and light peptides; (iii) averaged dot product values of ≥0.85 between library spectrum intensities and current observed intensities (dotp); (iv) averaged relative dot product values of ≥0.95 between the observed heavy and light ion intensities (rdotp); (v) coefficient of variation (CV) values of ≤20% for all dot product values; (vi) matching peak shape for precursor and product ions of both the heavy and light peptides. Individual peptide ions were quantified using the light-to-heavy area-under-the-curve ratio of the M, M+1 and M+2 peaks for MS1 and a manually curated set of *y-*ions for MS2. Curation was based on: (i) linearity as determined from the standard curve (**Supplementary Results, Supplementary Data 2**); and (ii) consistent observation between technical and biological replicates. To ensure only reliable ions were used for peptide quantification, further filtering was performed: (i) only light-to-heavy area-under-the-curve ratios with CV values of ≤20% between technical replicates were accepted; and (ii) the calculated value for each ion type had to fall within its pre-determined linear range. Finally, peptides were quantified by using the weighted average ratios between the M, M+1 and M+2 ions for quantification at the MS1 level and the selected *y-*ions for quantification at the MS2 level. This calculation was done in Skyline. All processed data related to the DIA-MS experiments can be found in **Supplementary Data 2**. All raw mass spectrometer and Skyline data files have also been deposited to the ProteomeXchange Consortium via the PRIDE partner repository (58) with the dataset identifier PXD010110.

### SEC-MALLS experiments and data analysis

Due to the instability and polydispersity of the NuDe and NuRD complexes upon concentration, size exclusion chromatography coupled to multiangle laser light scattering (SEC-MALLS) runs were carried out under denaturing conditions. NuDe and NuRD were first purified on GraFix sucrose gradients. The fractions containing the purified complexes were pooled, buffer exchanged (50 mM HEPES, 150 mM NaCl, 100 mM Tris, 6 M guanidine HCl, pH 8) and concentrated using Amicon^®^ Ultra, MWCO 100-kDa concentrators (Merck Millipore), pre-blocked with 5% (v/v) Tween-20. Chromatography runs were performed on a Superose™ 6 Increase 3.2/300 column (GE Healthcare) at 50 μL/min, in a buffer comprising 50 mM HEPES, 150 mM NaCl, 100 mM Tris, 2 M guanidine-HCl, pH 8. Similarly, both bovine thyroglobulin and bovine serum albumin (BSA) were purified by GraFix, and then subjected to SEC-MALLS under denaturing conditions. For untreated thyroglobulin and BSA samples, guanidine-HCl was omitted in all steps.

SEC-MALLS of the MTA^N^-HDAC-MBD_GATAD2CC_ complex was performed under non-denaturing conditions. The MTA^N^-HDAC-MBD_GATAD2CC_ sample used was eluted from FLAG-beads, after pulling down the complex from HEK cell lysate. Chromatography run was performed on a Superose® 12 10/300 GL (GE Healthcare) at 0.5 mL/min, in 10 mM HEPES, 75 mM NaCl, 0.2 mM DTT, pH 7.5.

All data analyses were performed using the ASTRA^®^ software (v6.1.1.17; Wyatt Technology). Both light scattering and UV data were used to calculate the molecular mass. Extinction coefficients (mL/(mg cm), at 280 nm) used were 1.148 for NuDe, 1.118 for NuRD, 1.044 for MTA^N^-HDAC-MBD_GATAD2CC_, 1.069 for bovine thyroglobulin and 0.614 for BSA. The BSA runs (untreated, 100 μL, 2 mg/mL) were used to align the UV and light scattering signals.

To calibrate the expected mass gain from glutaraldehyde crosslinking, we applied the following formula to the BSA and thyroglobulin standards: 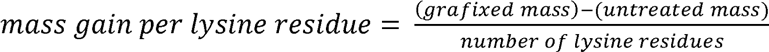 residues in BSA and thyroglobulin, respectively, we arrived at the conclusion that the GraFix protocol added an average of ∼540 Da per lysine residue (**Supplementary Results**).

### EM sample preparation

Post-sucrose density gradient separation, crosslinked fractions containing the complex of interest (based on the corresponding fractions collected from a non-crosslinked sucrose gradient run in parallel, **Figure S2c**) had the sucrose diluted to an approximate final concentration of 0.02% (w/v) by buffer exchange using an Amicon® Ultra, 100-kDa MWCO device (Merck Millipore), pre-blocked with 5% (v/v) Tween-20. The exchange buffer used was 10 mM HEPES-KOH pH 7.5, 75 mM NaCl, 0.3 mM DTT. The concentrated sample was centrifuged at 5,000 *g* for 5 min to remove possible aggregates and aliquots of the supernatant were frozen in liquid nitrogen and stored at –80 ^°^C. Final concentrations were in the range 3–30 ng/µL.

### Negative stain single particle electron microscopy

Sample (5 μL) was applied to a glow-discharged, carbon-coated 400-mesh copper grid (GSCu400CC—ProSciTech). After 2 min incubation time, the grid was blotted and washed with ten drops of distilled water, blotting on a filter paper in between washes. The grid was then washed in a drop of 1 % (w/v) uranyl acetate, blotted and subsequently incubated in another drop of 1% (w/v) uranyl acetate solution for 30 s, blotted and allowed to air dry at room temperature. Images were acquired using a Tecnai T12 TEM operated at 120 kV (MTA^N^-HDAC-MBD_GATAD2CC_, MTA-HDAC-RBBP and NuRD) or a Tecnai F30 TEM operated at 300 kV (NuDe), both equipped with a Direct Electron LC-1100 (4k × 4k) lens-coupled CCD camera. Images were recorded at a nominal magnification of 52,000×, 59,000×, or 67,000× corresponding to an unbinned pixel size of 2.79, 2.4 or 2.17 Å at the specimen level, respectively. Defocus values ranged from –1 to –2.5 μm.

### EM image processing and 3D reconstruction

All data processing was carried out using the SPHIRE (61) and RELION 2.0 software package (62). A total of 325, 469, 400 and 1332 micrographs were recorded for MTA^N^-HDAC-MBD_GATAD2CC_, MTA-HDAC-RBBP, NuDe and NuRD samples, respectively. For each data set, a subset of images was used for manual picking of around 1000 particles, subsequently used to generate templates for autopicking the complete data sets, with the exception of NuRD which was picked fully manually. The total number of extracted particles was 49216, 25155, 21622 and 124364 for MTA^N^-HDAC-MBD_GATAD2CC_, MTA-HDAC-RBBP, NuDe and NuRD, respectively. Various unsupervised 2D classifications were performed in order to select out poor quality particles (e.g., obvious aggregates, small particles).

For 3D classification and refinement of MTA^N^-HDAC-MBD_GATAD2CC_, reference-free 2D classes that had the highest distribution were selected for 3D classification. The 3D reference model used was a low pass filtered to 60 Å of the crystal structure of the MTA1_-_HDAC1 dimer (PDB: 4BKX). A single 3D class was produced containing 8233 particles, and final map was 3D refined and post-processed. The resolution of the final map was estimated at the post-processing stage using gold-standard Fourier shell correlation. All single particle image processing was performed with no symmetry assumed or imposed.

### Interaction studies using pulldown assays

Pulldown assays and western blot analyses used for the MTA^N^-HDAC-MBD_GATAD2CC_ subcomplex (in **Figure S5**) and PWWP2A interaction studies (in **Figure 5**) were performed essentially as described in (25) for proteins produced in HEK293 cells. Antibodies used for the western blots were from Cell Signalling Technology: anti-HA-HRP (2999S, 1:40,000) and anti-HDAC1-HRP (59581S, 1:80,000); from Sigma-Aldrich: anti-FLAG-HRP (A8592, 1:80,000); from Abcam: anti-MBD3 (ab157464, 1:2,500); from Leinco Technologies: anti-rabbit-HRP (R115, 1:10,000).

### Structural modeling

To model the MTA^N^-HDAC-MBD_GATAD2CC_ subcomplex, we combined the 3D structures available for NuRD subunits, XLMS data and our SPEM map. The structures used in our modeling were MTA1^ES^-HDAC1 (PDB: 4BKX), MBD domain of MBD3 (PDB: 2MB7) and MBD-GATAD2 coiled coil (PDB: 2L2L). For two NuRD regions, we constructed homology models using the SWISS-MODEL server (63). MTA1 BAH domain was based on the structure of the Sir3 BAH domain (PDB: 2FVU); and the MTA1 ZF domain was generated based on the C-terminal ZF of GATA1 (PDB: 2GAT). Finally, a single α-helix was built in PyMOL to represent the predicted helical region for MTA1(334-354).

Modeling was performed using HADDOCK 2.4 (48), these six described models and our XLMS data. Default run parameters were used with a high distance restrain of 5 Å between MTA-BAH(164) and MTA-ELM(165), as well as between MTA-SANT(333) and MTA-helix(334).

Resulting model was analyzed in Chimera (64), using the XLinkAnalyzer tool (65), and in PyMOL (The PyMOL Molecular Graphics System, Version 2.0 Schrödinger, LLC.). After building one copy of MTA-HDAC-MBD-GATAD2cc, we used two-fold symmetry to generate a second copy and build a model for the 2:2:2 MTA^N^-HDAC-MBD_GATAD2CC_ complex.

Haddock modelling of the MTA^C^-RBBP_2_ unit using previously published XLs (28) and XLs from this work was also performed. Based on existing structures of MTA^R1^-RBBP and MTA^R2^-RBBP complexes (**Figure S1b**), we extended the length of the MTA1 fragment in the latter structure *in silico* by drawing on sequence similarity to the longer MTA1^R1^. From the 71 XLs observed for the MTA^C^-RBBP_2_ unit, forty involved the region of MTA1 between R1 and R2 (residues 560–650), which is predicted to be disordered. Of the remaining XLs, three are unambiguous inter-subunit XLs between RBBP4 and RBBP7, demonstrating that ‘mixed’ complexes can form that contain more than one paralogue of a subunit. Although we had a model for the MTA^C^-RBBP_2_ unit from our previously published work (28), we performed new modeling here, taking into account the new MTA1^R1^-RBBP4 structure (30). No atomic model of MTA^C^-RBBP_2_ was able to fulfil all the XLs.

## Supporting information

Supplementary Results

## Acknowledgements

The work was funded by the following grants: National Health and Medical Research Council of Australia project grants: APP1012161, APP1063301, APP1126357 and a fellowship from the same organization to JPM (APP1058916). NIH R01-GM098264 to DCW. This research was facilitated by access to Sydney Mass Spectrometry, a core research facility at the University of Sydney

## Author Contributions

APGS, MST, MT, SRW and JWS prepared samples for EM experiments; APGS, SRW, JWS, LB and MJL contributed to EM data collection and processing; APGS, CS, JWS and JPM contributed to the modeling; JKKL, MST, MT and BLP contributed to DIA-MS and XLMS sample preparation and data collection; JKKL and BLP performed the DIA-MS and XLMS data analysis; MT and MS performed the PWWP2A experiments; SBH, DCW, GAB and NES provided intellectual input on the studies and manuscript; APGS, JKKL, MJL and JPM designed and supervised the studies, and wrote the manuscript with input from all the authors.

## Data availability statement

All mass spectrometry data that have been generated and analyzed during the current study have been deposited to the ProteomeXchange Consortium via the PRIDE partner repository. Accession numbers are PXD010110 for DIA-MS and PXD010111 for XLMS data. MTA^N^-HDAC-MBD_GATAD2CC_ EM map has been deposited in the EMDB repository with the accession numbers EMD-21382. These data are also available from the corresponding authors upon reasonable request.

---

**Figure S1.**
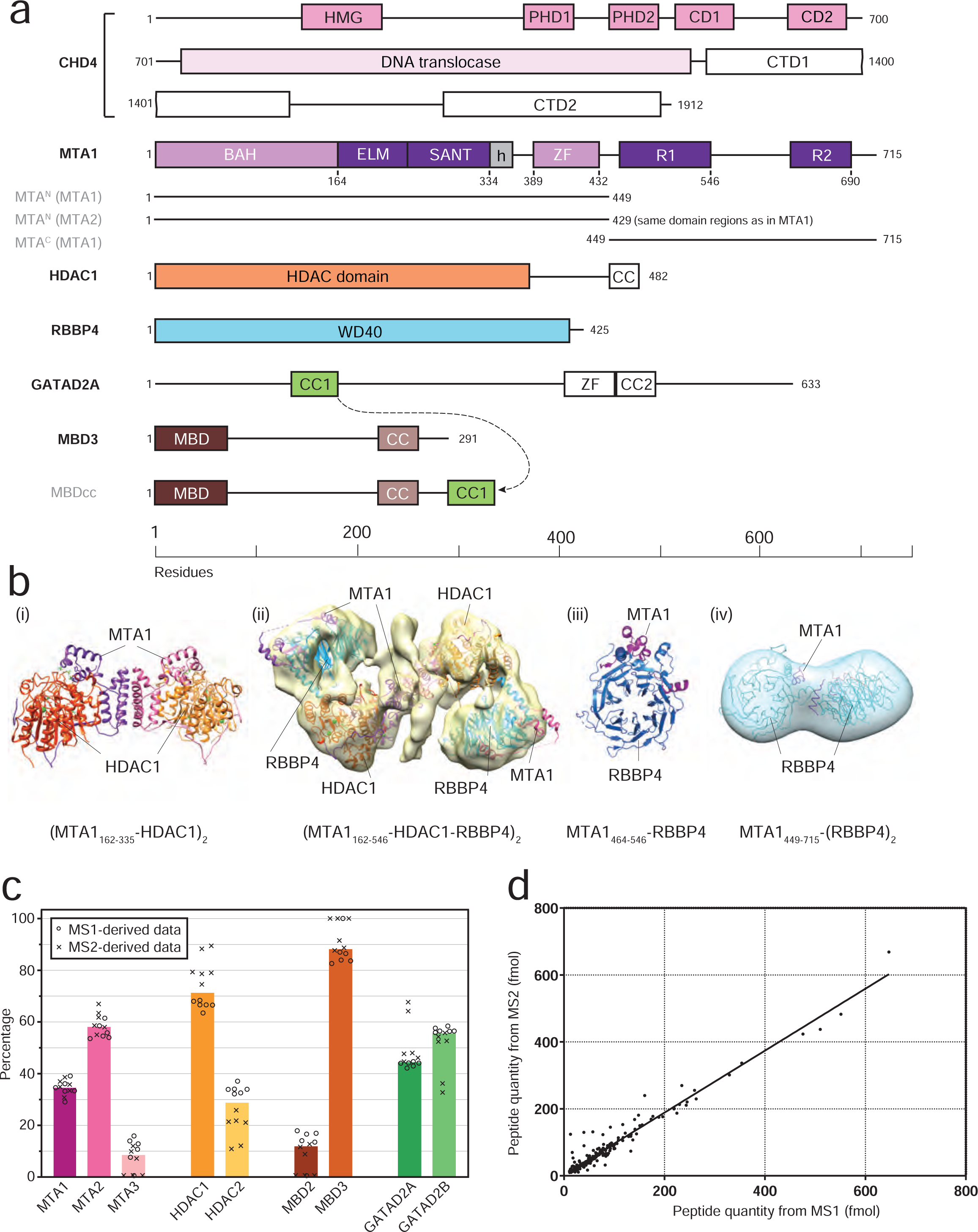
Composition of the NuRD complex and existing structural information, together with additional DIA-MS derived data. **a.** Domain structures of NuRD subunits. Numbering is shown for the human proteins. The six core NuRD components are labelled in black bold text and additional constructs used in this work are labelled in grey. MBD3cc is a fusion of MBD3 with the coiled-coil of GATAD2A, to which it is known to bind (46). Dark colours indicate domains for which structures are known; pale colours indicate domains for which reliable homology models can be built; white indicates regions predicted to be ordered but with unknown structures; and lines indicate regions predicted to be disordered. Paralogues of each subunit (*e.g.*, MTA2, HDAC2) generally have very similar domain structures. **b.** Structures of NuRD subcomplexes described in the text: (i) the MTA1_162–335_-HDAC1 complex (PDB: 4BKX, (27)) contains the MTA1 ELM and SANT domains; (ii) the MTA1_162–546_-HDAC1-RBBP4 complex (EMD-3399, (30)), contains the MTA1 ELM, SANT, zinc-finger and R1 domains; (iii) the MTA1_464–546_-RBBP4 complex (PDB: 5FXY(30)); and (iv) the MTA1_449–715_-RBBP4_2_ complex (EMD-3431, (28)). **c.** Relative abundance of subunit paralogues in the NuRD complex isolated from murine erythroleukemia cells and calculated DIA-MS data. Open circles and crosses indicate the individual data points, from each biological replicate, derived from MS1 and MS2, respectively. The bars show the median value. “n.d.” indicates a null value. **d.** Linear regression plot of the quantity of each NuRD peptide (across all samples – a total of 221 measurements) derived from MS1 measurements against values derived from MS2 measurements. The correlation coefficient was 0.97. The graph demonstrates that MS1 and MS2-based approaches are equivalent overall.

**Figure S2.**
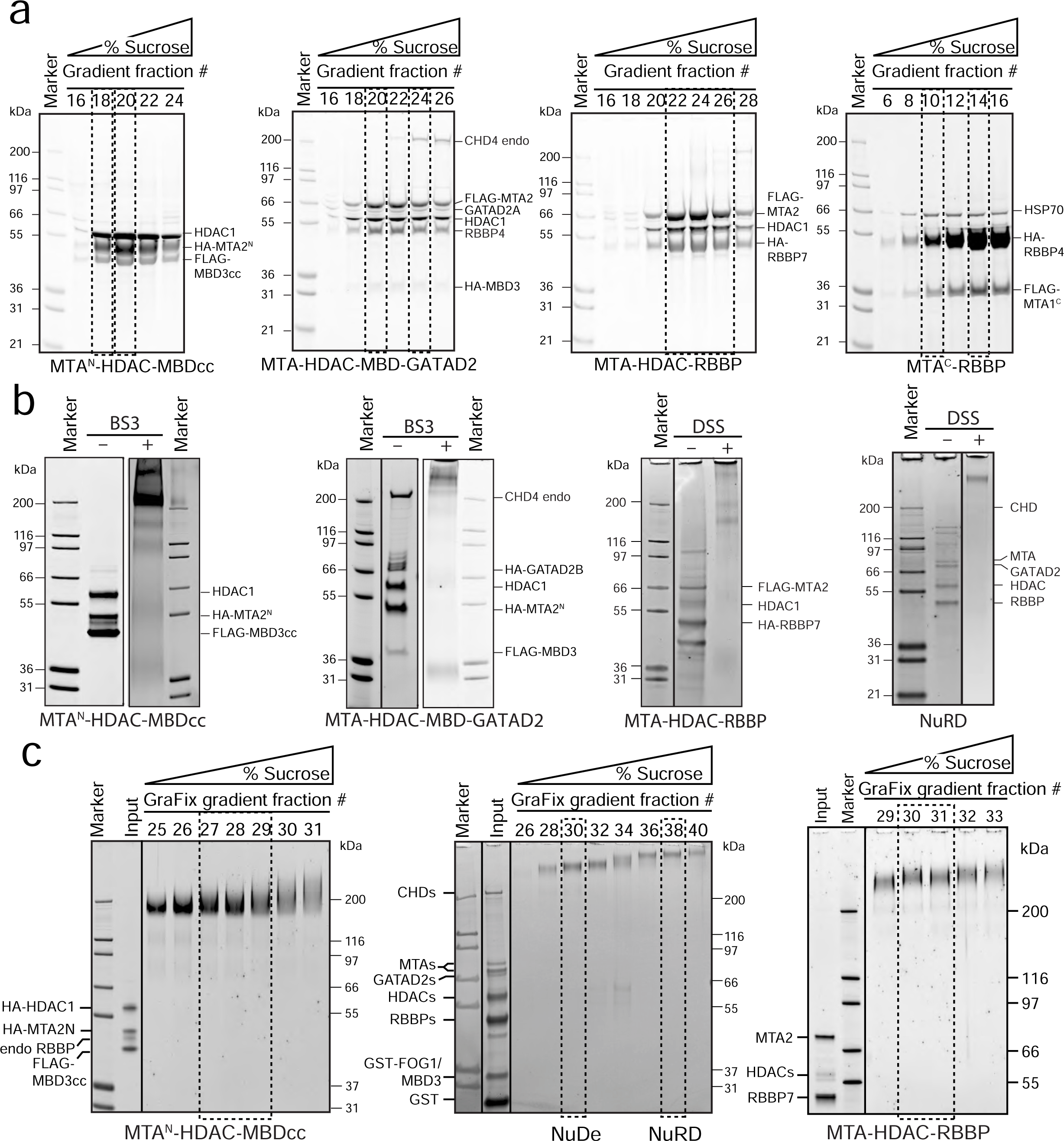
Purification of NuRD subcomplexes. SDS-PAGE showing purification of each of the complexes used for DIA-MS, XLMS or SPEM analysis. **a.** Fractions from sucrose gradient centrifugation for each complex. The fractions indicated with dashed boxes were used for DIA-MS measurements. Contaminating HSP70 was co-purified with MTA^C^-RBBP. **b.** Fractions from sucrose gradient centrifugation for each complex were selected, crosslinked with BS3, DSS or ADH (*data not shown*), and used for XLMS analyses. Solid lines indicate that lanes are from the same gel, in which some lanes have been cropped out. **c.** Fractions from sucrose gradient centrifugation for each complex – crosslinked with the GraFix protocol (55). The fractions indicated with dashed boxes were used for SPEM measurements. Solid lines indicate lanes that are from the same gel, but that some intervening lanes have been cropped out.

**Figure S3.**
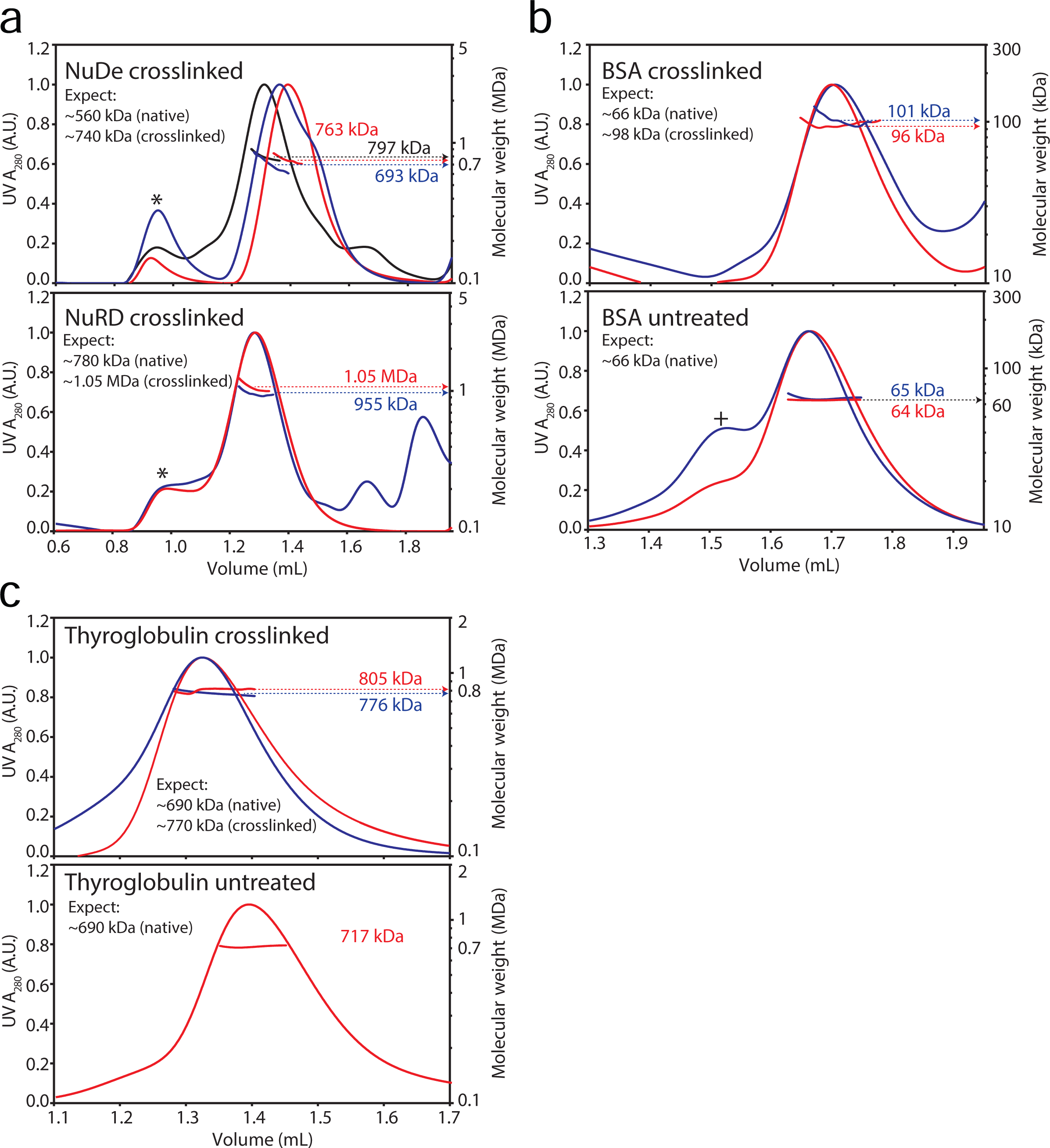
SEC-MALLS of NuRD and its subcomplexes. Replicate runs and calculated molecular masses are shown in different colours. **a.** Crosslinked NuDe and NuRD samples, which were run under denaturing conditions. “*” indicates high-molecular-weight aggregates (averaged measured mass > 8 MDa). **b.** Runs for crosslinked (top) and untreated (bottom) BSA. “+” indicates the position of a BSA dimer (measured mass = 120 kDa). **c.** Runs of crosslinked (top) and untreated (bottom) bovine thyroglobulin.

**Figure S4.**
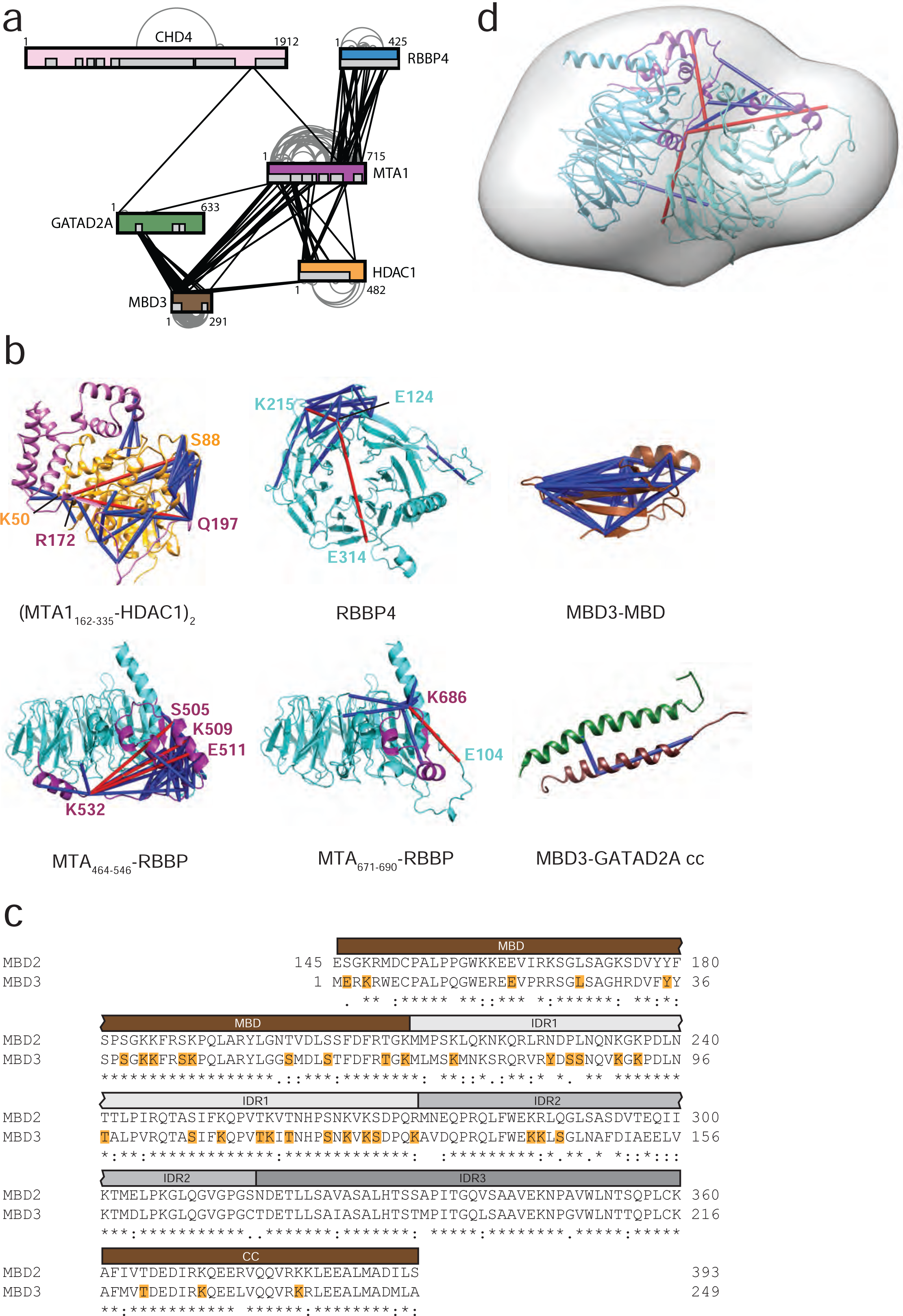
XLMS crosslinks are consistent with available structures. **a.** Compilation of unambiguous XLs observed in XLMS analysis of NuRD, NuDe, MTA-HDAC-RBBP, MTA^N^-HDAC-MBDcc and MTA-HDAC-MBD-GATAD2 complexes. Many of these XLs were observed multiple times within and between complexes. *Grey* boxes correspond to known and predicted protein domains. Inter-subunit crosslinks are in *black* while intra-subunit crosslinks are in *grey*. **b.** Observed XLs that can be mapped with existing structural data. 42 out of 44 observed XLs can be mapped onto the MTA1_162–335_-HDAC1 crystal structure (PDB: 4BKX); 20 out of 22 observed XLs can be mapped onto the RBBP crystal structure (PDB: 5FXY); all 41 observed XLs can be mapped onto the MBD3_1–68_ MBD domain (PDB: 2MB7). 36 out of 39 observed XLs can be mapped onto the RBBP4-MTA1_464–546_ structure (PDB: 5FXY). 7 out of 8 observed XLs can be mapped onto the RBBP-MTA1_671–690_ structure, for which the MTA1 fragment is based on PDB 4PBY, which was extended based on significant homology with the RBBP-MTA1_464–546_ structure. Both observed XLs can be mapped onto the MBD3_216–249_-GATAD2A_136–178_ coiled-coil structure (PDB: 2L2L). This region of MBD3 is highly conserved in MBD2. XLs not within its prescribed maximum length are shown in *red* and residues associated with these XLs are labelled. **c.** Crosslinked residues (*yellow*) within the MBD subunit, identified by XLMS. Only sequences from the MBD domain to the coiled-coil (CC) domain are shown. All crosslinks within the intrinsically disordered region (IDR) are localized to IDR1 and IDR2. Thus, 20/26 possible K/Y/S/T residues in IDR1 and IDR2 form XLs compared to 0/12 in IDR3.**d.** Ribbon representation of a molecular model of the MTA^C^-RBBP_2_ complex derived from XL-driven rigid-body docking in HADDOCK. The X-ray crystal structures of RBBP bound to C-terminal fragments of MTA1 (PDB:5FXY and EMD-3399) were used. The resulting model fits within our previously published low-resolution EM map of the MTA^C^-RBBP_2_ complex (EMD-3431), which is shown in *grey*.

**Figure S5.**
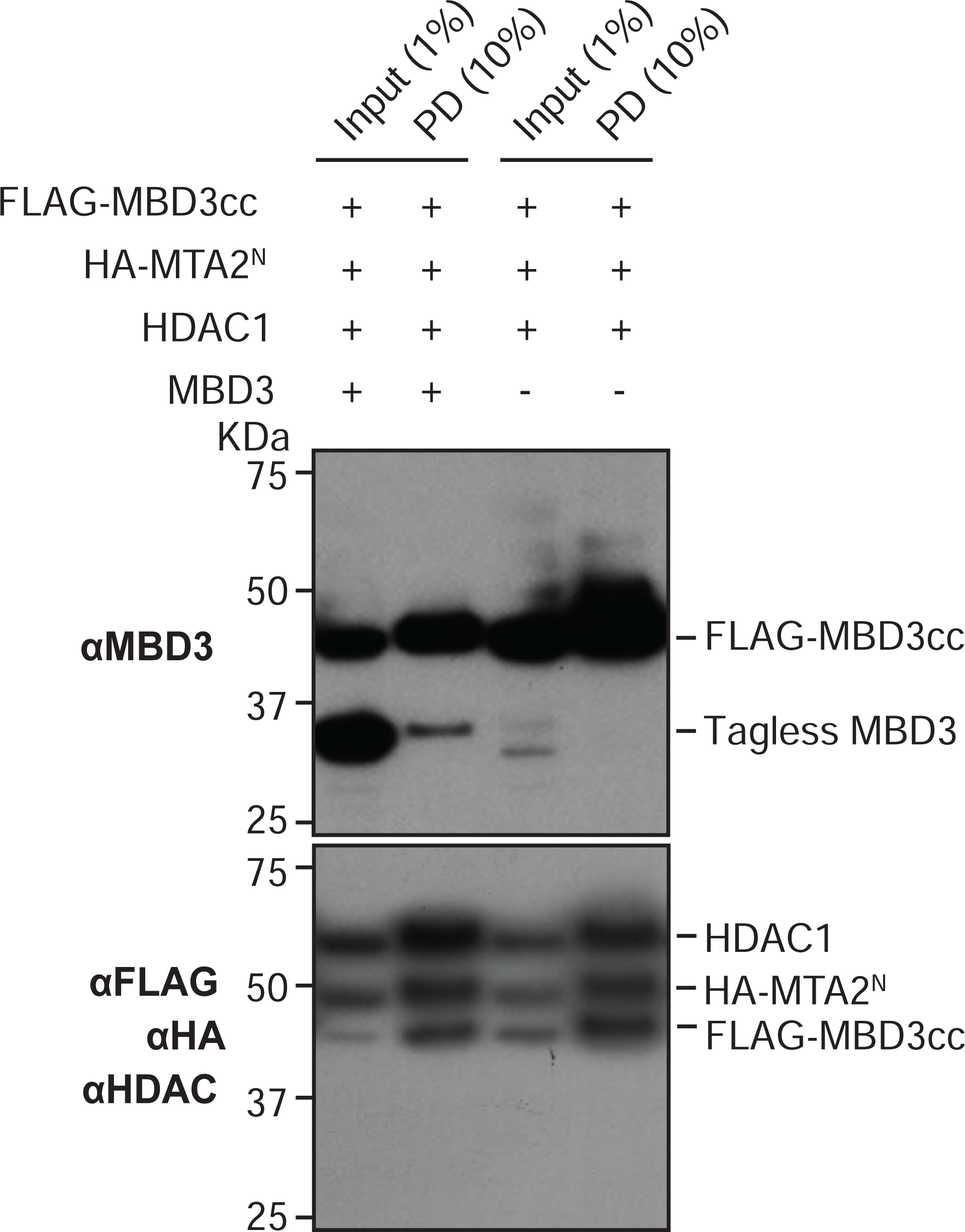
The MTA^N^-HDAC-MBD_GATAD2CC_ complex can contain more than one MBD protein. Western blot of input (lanes 1 and 3) and pulldown (lanes 2 and 4) samples from FLAG-affinity pulldowns in which FLAG-MBD3cc was co-expressed with HA-MTA2^N^, untagged HDAC1 and untagged MBD3 (lanes 1 and 2). Untagged MBD3 was not included in the control experiment (lanes 3 and 4). The top panel is an anti-MBD3 western blot, whereas the bottom panel was obtained by blotting against a mixture of FLAG, HA and HDAC antibodies.

**Figure S6.**
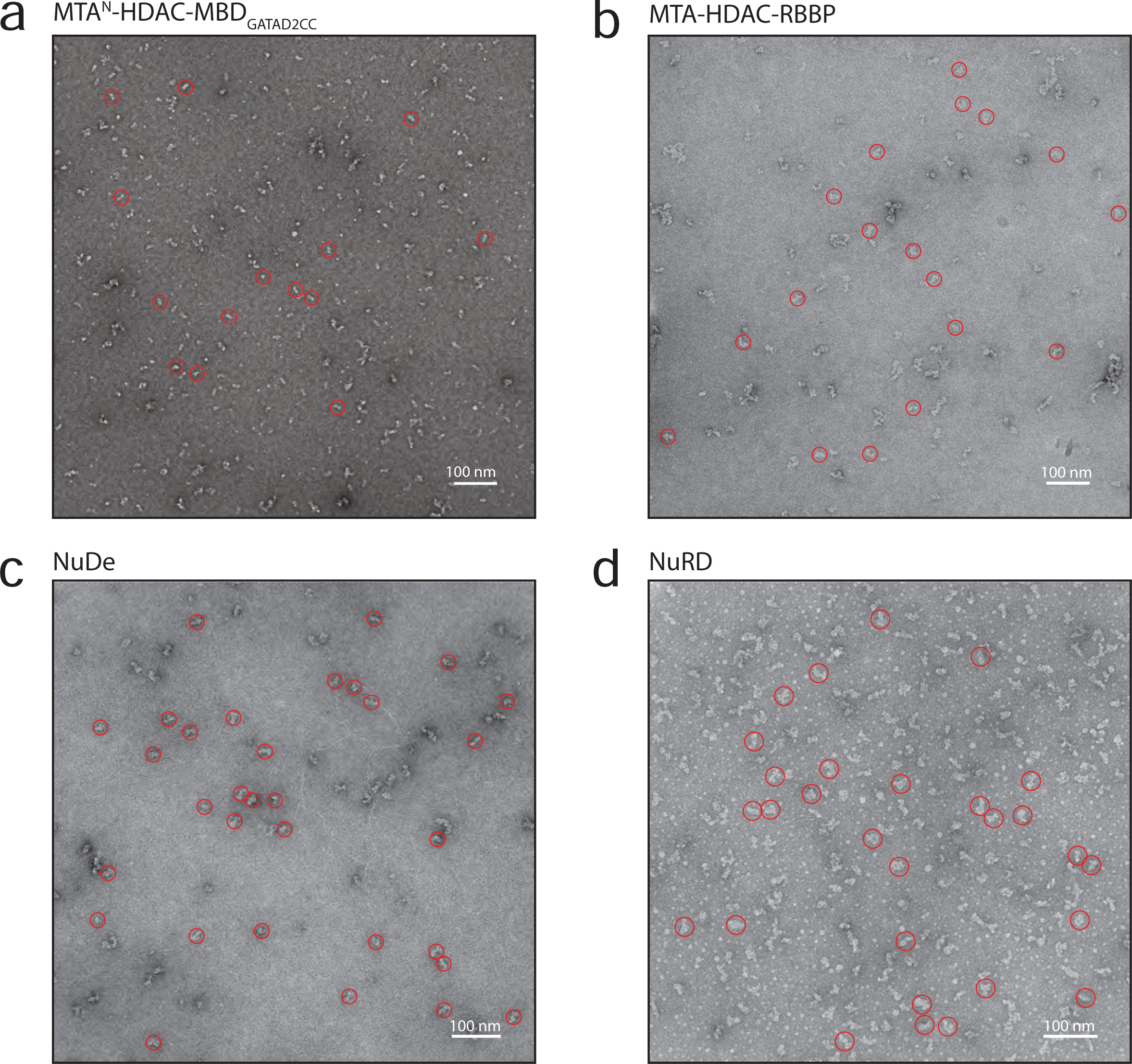
Representative negative-stained EM micrographs of NuRD complexes. **a.** MTA^N^-HDAC-MBD_GATAD2CC_ **b.** MTA-HDAC-RBBP. **c.** NuDe. **d.** NuRD. Representative particles are circled in *red*.

