## Supplementary Results for "The Nucleosome Remodeling and Deacetylase complex has an asymmetric, dynamic, and modular architecture"

**SUPPLEMENTARY RESULTS AND DISCUSSION**

***Peptide selection for MS-based quantification***

For quantification with internal standard peptides, the selection of reliable and representative proteotypic peptides (PTPs) for the protein targets of interest is an important step. Hence, in the first instance, we performed LC-MS/MS shotgun proteomics experiments to characterise our purified NuRD samples. From this proteomics dataset, we selected PTPs based on a pre-determined set of criteria: (1) Reproducible and reliable detection of the peptide across different MS runs and biological replicates; (2) Absence of missed cleavages; (3) Absence of known post-translational modifications (PTMs) (by querying protein databases including PhosphoELM (1) and UniProt (2) and large scale proteomic datasets that had not been incorporated into UniProt (at the time of experimental design) (3-5) or potential predicted PTM sites (GPS 2.1.1, (6)); (4) Hydrophobicity scores in the range 10¬46 (extremely hydrophobic peptides could present solubility issues and aberrant chromatographic behaviour); (5) Related to point (4), avoidance of series of hydrophobic residues (C, F, I, L, V, W, Y). A series of these residues can be difficult to synthesise; (6) Avoidance of extremely long or short peptides (9–10 residues is ideal); (7) Where possible, to avoid the following residues: methionine – unpredictable oxidation; cysteines – unpredictable oxidation but can be mitigated by reduction and alkylation of the samples; prolines – unusual elution profile due to isomerisation and predominant CID fragmentation pattern; N-terminal glutamines and glutamates – can undergo spontaneous modification to pyroglutamate; (8) Of less importance and where possible, to select for peptides with precursor ions <1000 *m/z* (this is due to the instrument used for data acquisition). Using these criteria, we selected 2–3 PTPs per target protein and where paralogues are very similar (*e.g.*, HDAC1/2 and RBBP4/7), an additional 1–3 ‘pseudo PTPs’ shared between the paralogues were also chosen. Selected peptides were synthesised with stable isotopes (carboxy-terminal Arg (^13^C_6_; ^15^N_4_) or Lys (^13^C_6_; ^15^N_2_)) as AQUA^TM^ peptides. A total of 31 peptides were chosen (exact numbers are indicated in parentheses): CHD4 (3), GATAD2A (3), GATAD2B (3) MTA1 (3), MTA2 (3), MTA3 (2), MBD2 (2), MBD3 (3), HDAC1 (2), HDAC2 (2), RBBP4 (3) RBBP7 (3). For HDAC1/2, one AQUA peptide was used for each HDAC1 and HDAC2, plus one additional ‘shared’ peptide for both HDAC1/2. For RBBP4/7, three ‘shared’ AQUA peptides were used. The list of peptides used is available in **Supplementary Data 2**. The linear intensity range of our selected peptides were determined as described below.

***Determination of linear intensity range of selected peptides***

We constructed a standard curve to assess the linearity of our heavy-labelled synthetic peptides in our MS quantification assay. We prepared and analyzed a dilution series of heavy-labelled synthetic peptides with a fixed quantity of highly purified NuRD proteins as the sample matrix. A range of 10–300 fmol was used a starting point, which was increased to 1000 fmol for the RBBP4/7 peptides. The MS data was processed in Skyline (7) as described in **Methods**. To determine the degree of linearity, we utilised simple linear regression analysis and response factor plots (8). From these experiments, 90 MS1 ions and 103 MS2 ions from 27 peptides (these numbers are doubled when considering heavy peptides as well) were found to have usable linear regions (**Supplementary Data 2**), with linear regression *R^2^* values of ≥0.972 and a response factor of <±20% of the average.

***MS1- and MS2-based quantification were highly correlated***

Recorded DIA-MS data were processed as described in the **Methods**. Excellent correlation was observed for quantification results between the MS1 and MS2 levels (*R^2^* = 0.95, Pearson *r* = 0.97, *n* = 221; **Figure S1d**); hence, we used both MS1- and MS2-derived data to inform us about the NuRD stoichiometry. All reported stoichiometry values in the text were median values from both MS1- and MS2-derived data. All processed data related to the DIA-MS experiments can be found in **Supplementary Data 2**.

***Detailed analysis of NuRD stoichiometry***

As we had multiple peptides to quantify each of our NuRD proteins, we were able to see that these peptides seldom agree with each other in terms of reporting the target protein’s quantity. There are a number of possible reasons why a particular peptide might under-report the true peptide quantity (*e.g.*, a protease-resistant endogenous peptide could lead to missed cleavages, or unknown PTMs on the endogenous peptide could alter the apparent peptide quantity). On the other hand, it was very unlikely that there would be instances of over-reporting. Thus, the commonly accepted solution of averaging multiple peptides for a target protein will result in under-reporting of the true quantity of the target protein. Hence, we opted to pick the highest quantified value for each protein instead – which should lead to a value that is as close to the true peptide quantity as possible.

To calculate NuRD stoichiometry, we first normalised all our quantified values to the average value between MTAs and HDACs ( $\frac{\left( MTA1+MTA2+MTA3 \right)+ \left( HDAC1+HDAC2 \right)}{2}$ ) then multiplied them by two. This yielded a value of ~2 for both MTAs and HDACs; it has been previously shown through X-ray crystallography that HDAC and MTA form a 2:2 complex (9) and subunit exchange data that show that the MTA and HDAC subunits are very stable in the NuRD complex (10). Based on these reasons, it was deemed reasonable to make both MTA and HDAC the stoichiometric reference subunits.

Our DIA-MS data also allowed us to interrogate the paralogue composition of the NuRD complex, which has not been examined in detail to date. **Figure S1c** show that, for the native complex isolated from MEL cells, each of the two GATAD2 paralogues are equally represented, whereas MTA2 (60%), HDAC1 (80%) and MBD3 (90%) are the predominant paralogues for those subunits. MTA3 and MBD2 are very low in abundance (≤10%). In all cases except for GATAD2, these percentages do not reflect the transcript abundance in MEL cells, suggesting that some selective incorporation of specific paralogues most likely takes place. Given that the CHD3/4/5 switch has clear functional consequences in neural development (11) and that MBD2 and MBD3 NuRD complexes can play distinct biological roles (reviewed in (12)), it is likely that shifts in the composition of each of the other subunit paralogues also translate to changes in activity. DIA-MS measurements made in different cell types in the future will be informative in this regard.

***Estimation of mass gain from glutaraldehyde crosslinking using SEC-MALLS***

As the NuRD and NuDe complexes were prone to aggregation upon concentration, we used NuDe and NuRD samples that had been purified using the GraFix protocol (13), which uses glutaraldehyde as the fixative agent. However, glutaraldehyde forms homopolymers with a range of sizes and the expected mass gain as a result of glutaraldehyde crosslinking is unknown. To this end, we first calibrated the expected mass gain from crosslinking by using two separate protein standards (BSA and thyroglobulin, **Figure S3b and c**). We concluded that the GraFix protocol adds ~540 Da to the mass of a protein per lysine residue (lysines are the major target of glutaraldehyde).

SEC-MALLS of the GraFix-treated NuRD complex yielded a mass of 980 ± 40 kDa (standard deviation from *n* = 2 measurements), which is within 7% of the predicted mass for the complex (accounting for crosslinking) of 1050 kDa (**Figure S3a**). For the NuDe complex, we measured a molecular mass of 760 ± 50 kDa (standard deviation from *n* = 3 measurements), which is within 3% of the predicted mass of 740 kDa (**Figure S3a**). The measured mass difference between the two complexes is also in very close agreement with the 218-kDa mass change expected from loss of a single CHD4 subunit.
